# A deep proteome and transcriptome abundance atlas of 29 healthy human tissues

**DOI:** 10.1101/357137

**Authors:** Dongxue Wang, Basak Eraslan, Thomas Wieland, Björn Hallström, Thomas Hopf, Daniel Paul Zolg, Jana Zecha, Anna Asplund, Li-hua Li, Chen Meng, Martin Frejno, Tobias Schmidt, Karsten Schnatbaum, Mathias Wilhelm, Frederik Ponten, Mathias Uhlen, Julien Gagneur, Hannes Hahne, Bernhard Kuster

**Affiliations:** Chair of Proteomics and Bioanalytics, Technische Universität München, Emil-Erlenmeyer-Forum 5, 85354 Freising, Germany; Computational Biology, Department of Informatics, Technical University of Munich, Boltzmannstr. 3, 85748, Garching bei München; Quantitative Biosciences Munich, Gene Center, Department of Biochemistry, Ludwig Maximilian Universität, 81377 München, Germany; OmicScouts GmbH, Lise-Meitner-Str. 30, 85354 Freising, Germany; Science for Life Laboratory, KTH - Royal Institute of Technology, Stockholm, Sweden; Department of Immunology, Genetics and Pathology, Science for Life Laboratory, Uppsala University, Uppsala, Sweden; JPT Peptide Technologies GmbH, Berlin, Germany; Center For Integrated Protein Science Munich (CIPSM), Munich, Germany

**Keywords:** Quantitative mass spectrometry, proteogenomics, human proteome, RNA-Seq, human transcriptome

## Abstract

Genome-, transcriptome- and proteome-wide measurements provide valuable insights into how biological systems are regulated. However, even fundamental aspects relating to which human proteins exist, where they are expressed and in which quantities are not fully understood. Therefore, we have generated a systematic, quantitative and deep proteome and transcriptome abundance atlas from 29 paired healthy human tissues from the Human Protein Atlas Project and representing human genes by 17,615 transcripts and 13,664 proteins. The analysis revealed that few proteins show truly tissue-specific expression, that vast differences between mRNA and protein quantities within and across tissues exist and that the expression levels of proteins are often more stable across tissues than those of transcripts. In addition, only ~2% of all exome and ~7% of all mRNA variants could be confidently detected at the protein level showing that proteogenomics remains challenging, requires rigorous validation using synthetic peptides and needs more sophisticated computational methods. Many uses of this resource can be envisaged ranging from the study of gene/protein expression regulation to protein biomarker specificity evaluation to name a few.

## Introduction

Delineating the factors that govern protein expression and activity in cells is among the most fundamental research topics in biology. Although the number of potential protein coding genes in the human genome is stabilizing at about 20,000, high-quality evidence for their physical existence has not yet been found for all and intense efforts are ongoing to identify these currently ~13% ‘missing proteins’ (Omenn et al. 2017). While it is also generally accepted that the quantities of proteins vary greatly within and across different cell types, tissues and body fluids (Wilhelm et al. 2014; Kim et al. 2014), this has not been quantified for many human tissues. Furthermore, it is not very clear yet how the many anabolic and catabolic processes are coordinated to give rise to the often vast differences in the levels of proteins. Messenger RNA levels are important determinants for protein abundance (Vogel et al. 2010; Schwanhäusser et al. 2011) and extensive mRNA expression maps of human cell types and tissues have been generated as proxies for estimating protein abundance (GTEx Consortium 2013; Uhlén et al. 2015; Thul et al. 2017). However, other studies have also highlighted the much higher dynamic range of protein abundance as well as rather poor correlation of mRNA and protein levels suggesting that further, and possibly diverse regulatory elements play important roles (Schwanhäusser et al. 2011; Liu, Beyer, and Aebersold 2016; Franks, Airoldi, and Slavov 2017). Decades of careful research has revealed numerous mRNA elements affecting translation or mRNA stability such as codon usage, start codon context or secondary structure to name a few. However, most of these studies focussed on single or few genes or single cell types or were performed in model organisms distinct from human systems and often did not cover a lot of proteins. Broader scale analyses have more recently become possible owing to advances in proteome and transcriptome profiling technologies, but these have mostly focussed on a single (disease) tissue or the cell-type resolved analysis of protein expression in single tissues (Zhang et al. 2014; Mertins et al. 2016). To the best of our knowledge, no broad-scale quantitative and integrative analysis of transcriptomes and proteomes across many healthy human tissues has been performed yet that would enable a comprehensive analysis of factors explaining the experimentally observed differences between mRNA and protein expression. Therefore, the purpose of this study was to generate a resource of molecular profiling data at the mRNA and protein level to facilitate the study of protein expression control and proteogenomics in humans. To this end, we analysed 29 major histologically healthy human tissues from the Protein Atlas Project (Uhlén et al. 2015) to provide a comprehensive baseline map of protein expression across the human body. As we show below as well as in an accompanying manuscript, this data can be used in many ways to explore protein expression and its regulation in humans. To facilitate further research on this fundamentally important topic and the many further uses that can be envisaged, all data is available in ArrayExpress (Kolesnikov et al. 2015) and proteomeXchange (Vizcaíno et al. 2014).

## Results and Discussion

### Comprehensive transcriptomic and proteomic analysis of 29 human tissues

We analysed 29 histologically healthy tissue specimen representing major human organs by label-free quantitative proteomics and RNA-Seq (Fig 1A). Tissues were collected by the Human Protein Atlas project (Fagerberg et al. 2014) and adjacent cryo-sections were used for paired (allele specific) transcriptome and proteome analysis. RNA-Seq profiling detected and quantified in total 17,615 protein coding genes with an average of 11,927 (+/− 937) genes per tissue (Fig 1B) when using a cut-off of 1 fragment per kilobase million (FPKM) (Uhlén et al. 2015). Proteomic profiling by mass spectrometry resulted in the identification and intensity-based absolute quantification (iBAQ) (Schwanhäusser et al. 2011) of a total of 15,210 protein groups with an average of 11,005 (+/−680) protein groups per tissue at a false discovery rate (FDR) of <1% at the protein, peptide, and peptide spectrum match (PSM) level (Fig EV1A). Protein identification was based on 277,698 non-redundant tryptic peptides, representing a total of 13,664 genes and, on average, 10,547 (+/− 512) genes per tissue covering, on average, 88% of the expressed genome in every tissue. While the total number of confidently identified proteins in this study is smaller than that of other (community-based) resources such as ProteomicsDB (Schmidt et al. 2018) and neXtProt (Gaudet et al. 2017) (coverage of 15,721 and 17,470 protein coding genes respectively), it provides a highly consistent collection of tissue proteomes including the deepest proteomes to date for many of the tissues analysed. It also provides protein level evidence for 72 proteins (represented by at least one unique peptide with Andromeda score of ≥100) that are not yet covered by neXtProt (release 2018-01-17; Table EV1).

**Figure 1.**
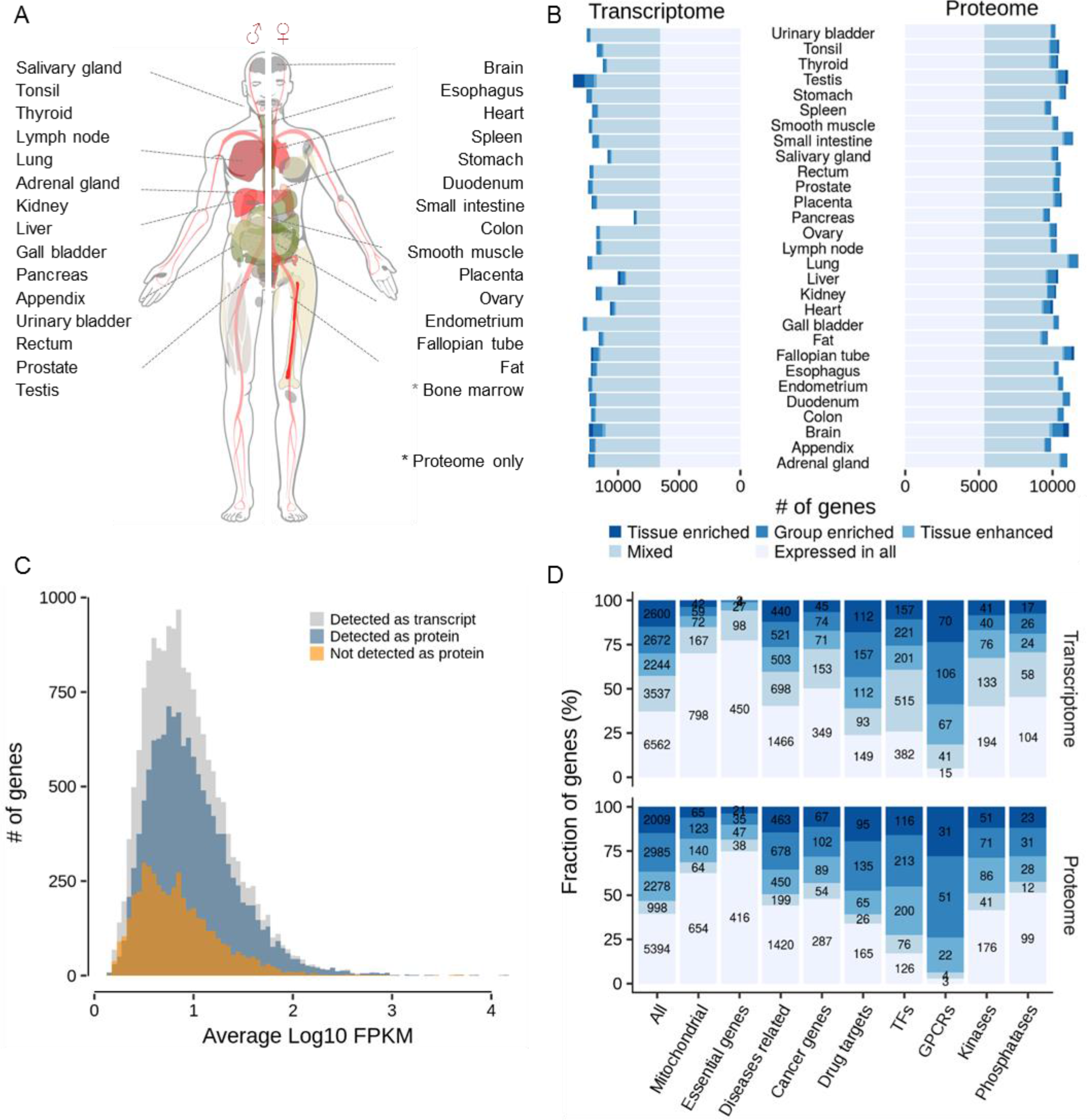
Comprehensive proteomic and transcriptomic analysis of 29 human tissues from healthy donors. (A) Body map of analysed tissues. (B) Number of genes detected on protein and mRNA level in each tissue. (C) Abundance distribution of all transcripts detected in all tissues (grey); the fraction of detected proteins is shown in blue and the fraction of transcripts for which no protein was detected is shown in orange. (D) Distribution of selected functional classes of transcripts and proteins across the expression categories shown in panel B. Colors are the same as in panel B.

Overall, 12,894 protein-coding genes were detected on both transcript and protein level, and the detected proteins spanned almost the entire range of mRNA expression again indicating very substantial coverage of the expressed proteome (Fig 1C). However, some proteins could not be detected even for highly expressed mRNAs (i.e. higher than the mean mRNA abundance). About 1/3 of these mRNAs were found in testis (486 of 1,574) and no other tissue contained nearly as many highly expressed mRNAs without protein evidence (Fig EV1B). The ‘missing’ proteins in the testis were statistically significantly enriched for processes related to spermatogenesis by gene ontology analysis (clusterProfiler; n=88 genes; BH adjusted p-value = 5×10^−12^). Although the rich expression of mRNAs in testis has been known for a long time and exploited for e. g. the cloning of many genes from cDNAs, the apparent absence of so many testis proteins with high mRNA expression is surprising. This was not due to e. g. poor coverage of the testis proteome (11,033 detected protein coding genes) or other obvious technical factors. Interestingly, almost 300 of these “missing” proteins have also not been detected by antibodies in testis (according to the Human Protein Atlas Project) and nearly 200 have no ascribed molecular function. The inability to detect these proteins by mass spectrometry or antibodies despite high levels of mRNA poses a number of questions. For example, are these proteins rapidly degraded implying specialized (and perhaps transient) functions in testis or sperm functionality? Are they perhaps stabilized in response to egg fertilisation? Proteins missing at the lower end of the mRNA expression range (less than mean mRNA abundance) are overrepresented in G-protein coupled receptor activity (n=170; BH adjusted p-value = 4×10^−45^), ion channels (n=111; BH adjusted p-value = 5×10^−7^)and cytokine related biology (n=121; BH adjusted p-value = 3×10^−10^). The abundance of these proteins may simply have been below the mass spectrometric detection limit or, as described many times, can be difficult to extract from cells owing to the presence of multi-pass transmembrane domains giving rise to few if any MS-compatible tryptic peptides after digestion. Interestingly, for 770 identified proteins, no corresponding mRNA was detected in any tissue (Fig EV1C, Table EV2). These proteins were enriched for e. g. immune related processes including Major Histocompatibility Complexes (MHC; n=40; BH adjusted p-value = 1×10^−41^) and antibodies (n=39; BH adjusted p-value = 2×10^−31^), that are either produced (on and off) by certain cell types in a given tissue or arise from elsewhere in the body not covered by our proteomes and transcriptomes.

To explore which and how many proteins show a tissue-specific expression profile, we applied the classification scheme of Uhlén et al. (Uhlén et al. 2015, 2016) previously developed for mRNA profiling and which stratifies genes into the five classes “tissue-enriched” (5-fold above any other tissue), “group enriched” (5-fold above any group of 2-7 tissues), “enhanced” (5-fold above the average of all other tissues), “expressed in all” (expressed in all tissues) as well as “mixed” genes (which do not match the other categories). Overall, a large fraction of all represented genes was expressed in all tissues: 37% (6,562) at the transcript level and 39% (5,394) at the protein level. However, 43% (7,516) of all transcripts and 53% (7,272) of all proteins showed elevated expression in one or more tissues (“tissue-enriched”, “group-enriched” or “tissue-enhanced”). Only 4.3% (on average) of all transcripts and 5.4% of all proteins showed a tissue-enriched profile. Two notable exceptions are brain and testis which exhibit a higher percentage of tissue enriched proteins and transcripts in line with a recent analysis of RNA-Seq data from the Human Protein Atlas and GTEx projects (GTEx Consortium 2013). Proteins with more tissue restricted expression tended to be of somewhat lower abundance (Fig EV1D).

The above global trends in transcript and protein tissue expression distributions were also mirrored by functional categories of genes but with some interesting detail (Fig 1D). For example, while the tissue distribution of expression of disease-associated genes followed that of all genes, the expression of drug targets in general and GPCRs in particular was much more tissue restricted speaking to the notion that proteins may make for better drug targets if they are not ubiquitously expressed (Hao and Tatonetti 2016). In this context, we point out that our baseline map of protein expression across the human body may be of general value to drug discovery as one can e. g. quickly examine the expression of a particular target of interest, to help to better understand adverse clinical effects and off-target mechanisms of action of drugs based on their tissue expression profiles. For instance, a recent study revealed phenylalanine hydroxylase (PAH) as an off-target of the pan-HDAC inhibitor panobinostat (Becher et al. 2016). Our map of protein expression shows that PAH is abundantly expressed in liver (and kidney) which is also the major site of hydroxylation in the human body (Matthews 2007), indicating that the liver is the major site where panobinostat exerts its detrimental effects, i. e. leading to decreased tyrosine levels, and eventually hypothyroidism in affected patients. In contrast, essential genes (Wang et al. 2015; Hart et al. 2015; Blomen et al. 2015) as well as mitochondrial genes were found in the vast majority of all tissues in line with their central roles for maintaining cellular homeostasis. Despite the differences in detail, our dataset confirms, at the protein level, that there is a core set of ubiquitously expressed genes/proteins and that individual tissues are not strongly characterized by the categorical presence or absence of mRNAs or proteins but rather by quantitative differences (Geiger et al. 2013). This is also evident from an analysis of the most divergently expressed proteins or transcripts that shows enrichment of proteins related to the functional specialization of the respective tissue (Fig EV1E, Table EV2).

### mRNA and protein expression

The dynamic range of transcripts detected by RNA-Seq spanned about four orders of magnitude and that of proteins detected by mass spectrometry spanned eight orders of magnitude (Fig 2A). This difference alone explains (at least in part) the overall higher coverage of the expressed proteome by RNA-Seq compared to that of LC-MS/MS. The much wider dynamic range at the protein level implies that protein synthesis and protein stability play an important role in determining protein levels beyond mRNA levels. Moreover, the number of protein copies produced per molecule of mRNA appears to be much larger for high- than for low-abundance transcripts, leading to a nearly quadratic relationship between mRNA levels and protein levels in every tissue (slope of 2.6 in Fig 2B (brain) and between 1.8 and 2.7 for all 29 tissues, Fig EV2A; Appendix Fig S1). This may be rationalized by cellular economics such that genes encoding highly abundant proteins not only express high mRNAs levels, but also encode regulatory elements that favour high translation efficiency and high protein stability (Vogel et al. 2010). The often vast differences in mRNA and protein expression within a tissue can also be visualized by plotting the ranked order of relative intensities of transcripts and proteins (Fig 2C). For example, in the heart, 41% of the total mRNA quantity (by FPKM) represents a single protein (MT-ATP8) and nearly 60% of all mRNA covers just five transcripts (all coding for mitochondrial proteins). In contrast, about 13% of the total protein quantity (by iBAQ) is contributed by five proteins (four of which are myosins and one represents a ‘contamination’ from blood present in the tissue). One would expect the heart to be rich in both protein families owing to the contractile function of the organ which requires a lot of energy. And while it is possible that mitochondrial proteins are underrepresented in quantitative terms (they are not underrepresented merely by counting presence/absence) because our lysis conditions may not have solubilized this organelle with high efficiency, it is surprising that even among the 100 most highly expressed mRNAs and proteins, only about 20% are the same (Fig 2D) and the overlap only increases to about 60% for the 5,000 most abundant proteins and transcripts (Fig EV2B). The abundance distribution of transcripts and proteins is also quite different between tissues with the spleen showing the opposite characteristics compared to the heart, and the lung showing a more even distribution of transcript and protein levels (Fig EV2C-D). Due to the fact that a majority of the proteins are expressed at similar levels across human tissue, it is not very surprising that the correlation of mRNA/protein ratios across tissues is generally not very strong (Fig 2E; median 0.35). Still, there is positive correlation in almost 90% of all cases and almost half are also statistically significant. However, care has to be taken when interpreting this distribution. We generally find that proteins and transcripts that are high (low) expressed in one tissue are also expressed high (low) in many (but not always all) other tissues (Fig EV2E). As shown in Fig 2F, the transcript and protein levels of the tyrosine kinase SYK are highly correlated across tissues reflecting the specialized function of the protein in T- and B-cell biology. In contrast, other proteins such as EIF4A3 (a DEAD-box RNA helicase involved in translation initiation) show no such correlation. However, this is merely the result of similar expression levels in most tissues reflecting also their roles in central biological processes in all tissues (Wilhelm et al. 2017).

**Figure 2.**
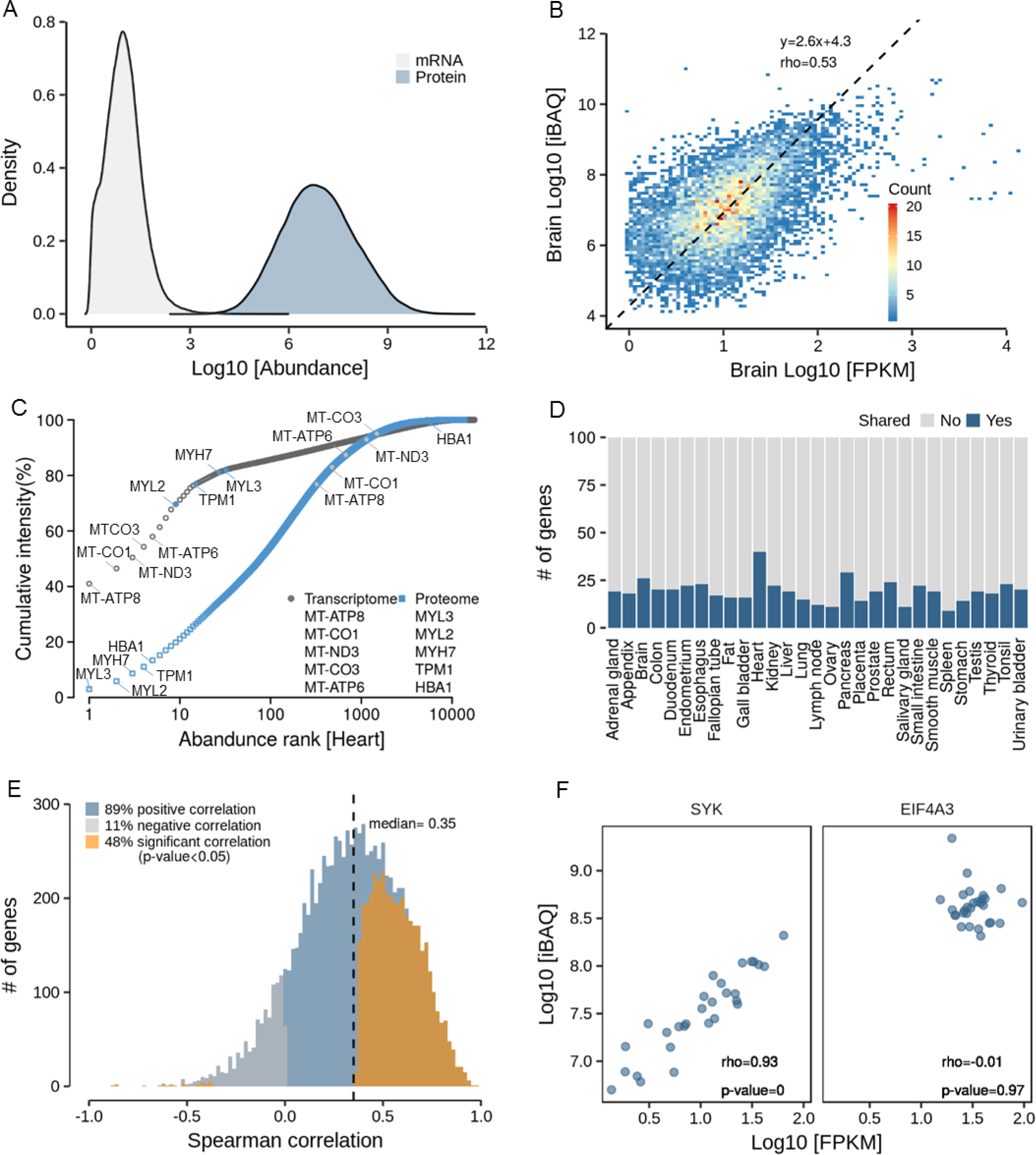
Analysis of protein and transcript expression levels within and across tissues. (A) Distribution of global transcript and protein abundance in all tissues. It is apparent that the dynamic range of protein expression vastly exceeds that of mRNA expression. (B) Protein to mRNA abundance plot for brain tissue. The slope of the regression line indicates that high abundance mRNAs give rise to more protein copies per mRNA than low abundance mRNAs. (C) Ranked abundance plot of proteins and transcripts in human heart. While the 10 most abundant transcripts cover almost 70% of all transcripts in this tissue, the corresponding proteins only represent about 20% of the total protein. (D) Analysis of the number of genes that are shared among the 100 most abundant transcripts and proteins. Regardless of the tissue, the fraction of shared genes rarely exceeds 20%. (E) Correlation analysis of the expression of proteins across all tissues. Almost 90% of all proteins show a positive correlation across tissues. (F) Examples for proteins that show high (SYK, left panel) or no (EIF4A3, right panel) correlation of protein expression across tissues. While the former indicates that different tissues express different quantities of SYK, EIF4A3 expression appears to be similar in all tissues.

It is noteworthy that proteomes correlate stronger between tissues (median of 0.77) than transcriptomes (median of 0.67) (Fig 3A). This might be due to the fact that the dynamic range of protein levels is larger and thus small biological or technical variations of individual genes have a negligible impact on the overall rankings. It might also imply that there are mechanisms in cells that buffer the protein quantities against changes in mRNA abundance (Liu, Beyer, and Aebersold 2016; Kustatscher, Grabowski, and Rappsilber 2017). The strongest correlations both for transcripts and proteins were found for the anatomically adjacent small intestine and duodenum. At the proteome level, the brain and heart show clear differences to other proteomes and gastrointestinal organs appear to be more similar to each other. Visualizing the transcriptome and proteome profiles in a plane using co-inertia analysis (CIA) (Culhane et al. 2005) indicate that mRNA and protein levels are more similar to each other within tissues than between tissues (Fig 3B) also reflected by an RV coefficient of 0.78 (a multivariate generalization of the squared Pearson correlation coefficient). Moreover, the CIA grouped several tissues according to similarities in their physiological function with tissues of the immune system and of the gastrointestinal tract representing the largest groups. It is interesting to note that this clustering appears to be driven by the cellular composition of individual tissues. For instance, the appendix co-clusters with the spleen, lymph node and tonsil and all four tissues contain a large fraction of lymphocytes (Fig 3C, blue panel). Similarly, the stomach, duodenum, small intestine, colon and rectum all comprise a large proportion of (intestinal) glandular cells, which are important determinants of the molecular make-up of those tissues (Fig 3C, grey panel). All the above illustrates that there must be multiple molecular factors and mechanisms determining the quantitative expression of protein. This particular aspect of the present mRNA/protein expression resource may be particularly useful for the community as it provides a rich data source for the study of protein expression control (see also the accompanying manuscript).

**Figure 3.**
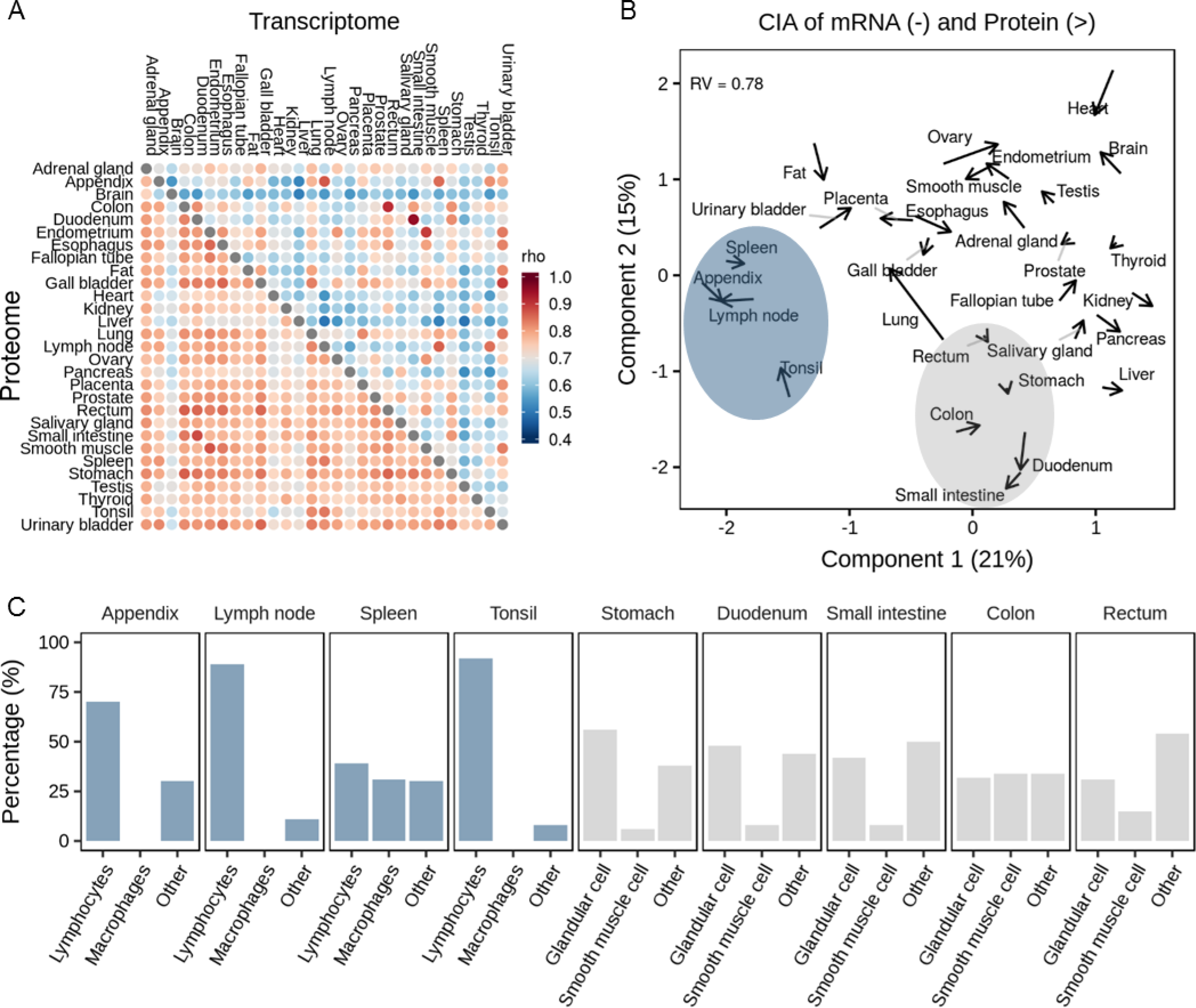
Correlation analysis of protein and transcript expression levels. (A) Global correlation analysis of proteomes and transcriptomes across human tissues. It is apparent that proteomes correlate stronger between tissues than transcriptomes. (B) Co-inertia analysis of transcriptome and proteome levels of all 29 tissues (arrow base: transcriptome; arrow head: proteome) showing that some tissues show similarities in transcript and protein expression profiles. (C) Average cellular compositions of selected tissues showing that the similarities found in panel B are largely driven by similarities in cell types.

### Proteogenomic characterisation of human tissues

One aspect of the data we cover in more detail in this study is the considerable interest in the community to use proteomics data to annotate genomes, often referred to as proteogenomics. With matched RNA-Seq and proteomics data at hand, we set out to assess the merits of proteogenomics at several levels. First, we investigated the identification of protein isoforms. Based on RNA-Seq data, it has been suggested that human cell types typically express one dominant isoform (Ezkurdia et al. 2015). In proteomics, isoforms are much more difficult to distinguish unambiguously because the identification of proteins is inferred from the underlying peptide data. Given that many proteins contain conserved short stretches of amino acids and the fact that the median sequence coverage achieved for each protein is limited (here between 14 and 25%; Fig EV3A), many potential isoforms will not be covered by unique peptides and some peptides may also match to multiple entries in comprehensive sequence collections such as Ensembl (102,450 entries), thus often identifying a so-called protein group rather than one specific protein or isoform thereof. Illustrated by the proteomic data obtained from trypsin digested tonsil (Fig 4A), only 14% of all protein groups contained one single protein when searching the MS data against Ensembl. However, when searching the same data against a protein sequence database constructed from the tissue specific RNA-Seq data, the proportion of single entry protein groups increased to 53% (see Appendix Fig S3 for all tissues). In this way, we were able to identify 53,858 non-redundant isoforms by RNA-Seq for 17,560 genes and confirm 15,257 by proteomics for 11,833 genes across the 29 tissues (Fig EV3B, Table EV1 and EV6).

**Figure 4.**
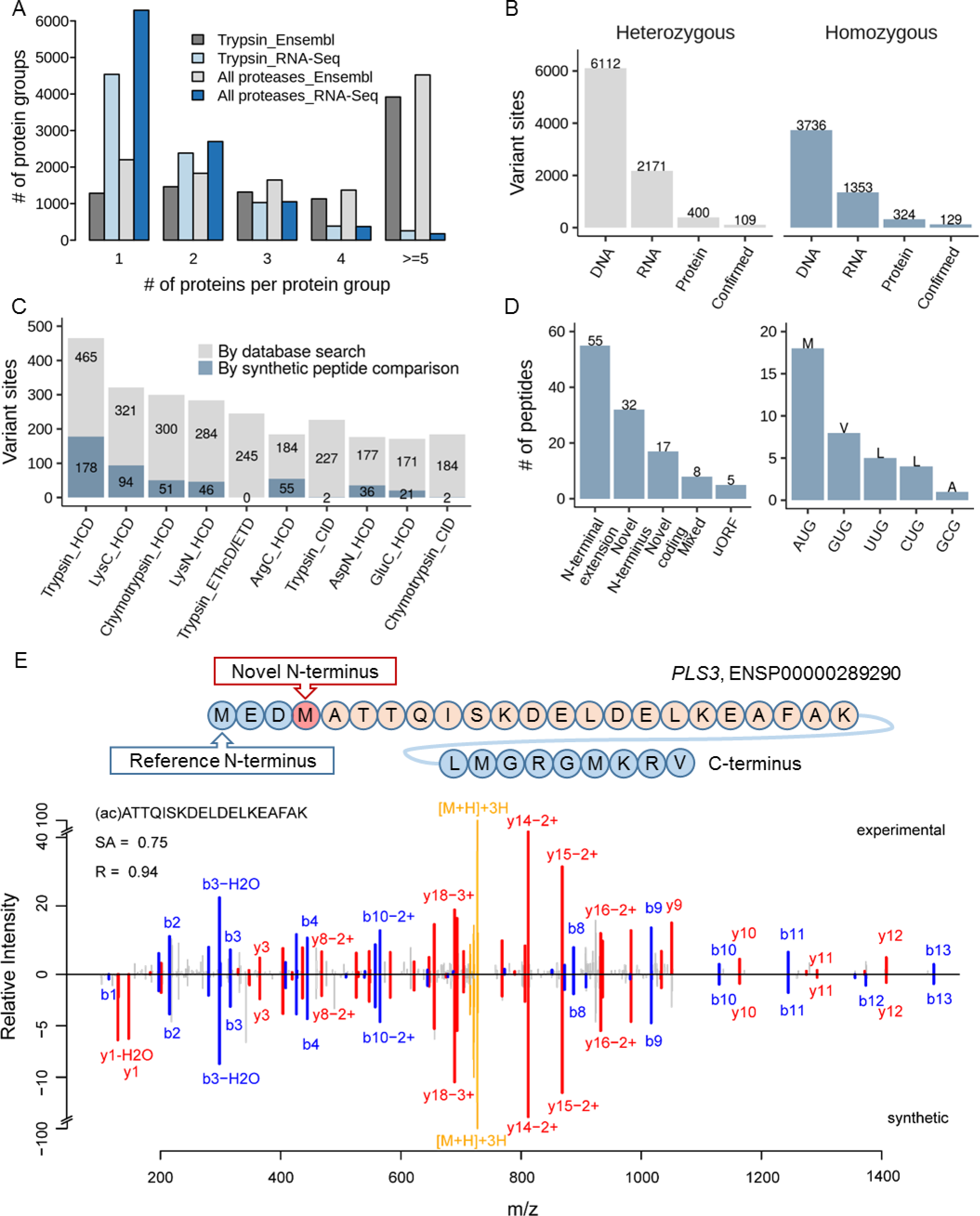
Proteogenomics exploration for protein-level detection of isoforms, single amino acid variants and alternative translation sites. (A) Searching the tonsil proteomic data (trypsin alone or all enzymes) against a tissue-specific sequence database constructed from RNA-Seq data drastically reduces the number of individual protein sequences in protein groups compared to searches against Ensembl, allowing for the more efficient detection of protein isoforms. (B) Number of single amino acid variants detected by whole exome sequences, RNA-Seq, mass spectrometry and validation using synthetic peptide spectra comparisons. It is apparent that only a very small fraction of all variants detected at the DNA or RNA level can be detected at the proteome level. (C) Analysis of which proteomic workflow contributed to the confident detection of single amino acid variants. (D) Results of the detection of non-canonical coding regions using proteomics data (left panel) and the alternative start codon identified by acetylated N-terminal peptides (right panel). The majority of cases are N-terminal extensions of annotated genes. All but one of the detected alternative translation start sites correspond to point mutations of the first base of the classical AUG codon. (E) Validation of a novel translation start site for the protein PLS3. The upper panel shows the novel translation site position within the amino acid sequence context and the lower panel shows the tandem mass spectra of the endogenous N-terminally acetylated peptide and the corresponding synthetic peptide spectrum.

One way to improve the detection of isoforms is to increase the sequence coverage in the proteomic data. To this end, we performed an ultra-deep proteomic analysis of tonsil tissue by applying seven different proteases (trypsin, LysC, LysN, GluC, ArgC, AspN, and chymotrypsin) and three peptide fragmentation techniques (HCD, CID and EThcD/ETD). This resulted in the identification of 11,569 protein groups (10,288 genes), represented by 421,073 non-redundant peptides leading to a median protein sequence coverage of 54% (Fig EV3C-E; Table EV5; when searched against Ensembl). Of these protein groups, 2,201 could be unambiguously linked to a single isoform. When searched against protein sequences derived from the tonsil-specific RNA-Seq data, we identified 10,592 protein groups, among which 6,293 represented a particular isoform identified by unique peptides (Fig 4A). The above shows that isoform calling on the protein level is possible, but doing so systematically remains challenging. We further note that because most isoforms were detected by very few isoform-unique peptides, confident quantification of the different isoforms of the same gene found in the same tissue is currently very difficult and may require targeted MS assays rather than shotgun approaches. In this context, it is also worth mentioning that there is no clear consensus in the proteomics and transcriptomics communities as to how quantitative values should be allocated to particular proteins or transcripts. While it is custom in proteomics to use the parsimonious approach (i. e. allocate all iBAQ intensity to the protein with the highest overall evidence), it is custom to distribute RNA-Seq reads covering shared sequences across multiple transcripts containing that sequence (Trapnell et al. 2010). It would not be surprising if these differences in quantification approaches would add substantially to the poor correlation of mRNA and protein levels (or their ratios). In addition, there is currently no tractable way to determine which allele of a gene gave rise to a detected protein or isoform thereof.

To assess the ability of proteomics to detect genetic variants, we analysed whole exome sequencing (WES), RNA-Seq and ultra-deep proteomics data of tonsil tissue. In the WES data, the average exon coverage was 98x and 97% of the exons were covered >20x providing a sound basis for the identification of single amino acid variants (SAAVs). Variant calling and filtering of WES data resulted in 9,848 high-quality, non-synonymous point mutations (i.e. nonsense and missense variants excluding I>L and L>I variants that cannot be distinguished by mass spectrometry), representing 5,527 human genes and including 6,112 heterozygous and 3,736 homozygous variants (Fig 4B, Table EV6). In the RNA-Seq data, 3,524 of the 9,848 genomic variants (36%; 2,171 heterozygous and 1,353 homozygous cases; representing 2,428 genes) were covered sufficiently (≥10x) to assess their genotype. The reason for the substantial loss of coverage in the RNA-Seq vs exome data is because i) not all genes are expressed from both alleles in a given tissue and ii) even at a sequencing depth of 50 million reads, the dynamic range of mRNA abundance is too high to cover all transcripts and variants many times over. It has been noted before that the identification of SAAVs by proteomics is challenging and plagued by false positives in standard database searching regimen because i) the tandem mass spectra used for database searching are often not of very high quality, ii) these spectra often do not contain the complete amino acid sequence information of the underlying peptide and iii) the current FDR statistics for peptide/protein identification do not translate well to variant calling on the peptide level. As a result, random matches can and will frequently occur raising substantial concern about part of the current proteogenomic literature (Nesvizhskii 2014). When searching our proteomic data against concatenated sequences obtained from the WES data, Ensemble and UniProt and requiring an identification by both Mascot and Andromeda as well as a number of further criteria (for details see methods), we identified 1,942 candidate peptides mapping to 724 of the 9,848 (non-canonical) exome variants (7.4%; 400 heterozygous and 324 homozygous cases). These peptide variants are all missense mutations (Table EV6). For 41% of the heterozygous cases (165 out of 400), we obtained peptide level evidence for the canonical and alternative variant, while for the remaining cases, we only identified the alternative variant (235). For validation, candidate peptide spectra were compared to those of synthetic reference standards (Zolg et al. 2017). To this end, we attempted the synthesis of reference peptides for all 724 alternative variants and obtained such peptides for 574 cases. Automated spectral angle analysis (Toprak et al. 2014) provided evidence for 238 variants (SA ≥0.7, Mascot ion score ≥50) including 109 heterozygous and 129 homozygous cases (Fig 4B; Table EV6). Manual inspection of the above 724 candidate peptides identified 204 unique alternative variant sites of which 158 were also found in the SA analysis. The variants that passed our (conservative) filtering criteria merely represent 2.4% of all variants detected at the exome level, 6.7% of the variants detected at the mRNA level and 32% of the candidates suggested by database searching. When tracing the confidently identified peptide variants back to the proteomic workflow, it became clear that the vast majority of all variants are represented by peptides generated by Trypsin, LysC and ArgC cleavage and using the standard HCD fragmentation technique. In addition, the confirmation rate (using synthetic peptide reference standards,) for tryptic peptides was also much higher than that of the other enzymes (Fig 4C; synthetic peptides were not measured by EThcD/ETD). This can be attributed to the fact that trypsin-like peptides generally show well predictable fragmentation behaviour and that most bioinformatic tools are optimized for use with data generated from tryptic digestion of proteomes. While the above shows that some of the variants detected on the nucleotide level could be confirmed at the protein level, the overall success rate was low. We note here that this was not due to lack of expression of the underlying gene because the proteomic data covers 76 % of all expressed tonsil genes (8,869 of 11,746 mRNA-Seq genes), 47% (2,615 of 5,527) of all the genes for which variants were detected by exome sequencing and 75% (1,822 of 2,428) by RNA-Seq respectively (further discussed below). Instead, the main reasons for poor coverage of variants at the proteome level are the still limited sensitivity and dynamic range of detection, limited peptide coverage of a protein and the insufficient coverage of amino acids in peptide mass spectra along with shortcomings in peptide identification algorithms.

Recent research showed that there is more heterogeneity in gene models than previously anticipated, as a result of e. g. alternative translation initiation sites (aTIS) (Na et al. 2018) and there is an on-going debate in the community whether or not long non-coding RNAs (lncRNAs) can be translated into proteins (Chen et al. 2017). Ribosomal profiling showed that thousands of potential aTIS may exist and that ~40% of all lncRNAs can at least engage the ribosome (Kearse and Wilusz 2017). In order to explore if our resource can provide protein evidence for such cases, we used a database search strategy (Marx et al. 2017) in which we searched all LC-MS/MS files against combined sequences from i) a curated lncRNA database (GENCODE v.25), ii) a database containing protein sequences derived from alternative translation initiation sites (see methods), iii) GENCODE, iv) UniProt and v) the tissue-specific RNA-Seq data. Any potential lncRNA or aTIS peptide was required to originate from just one single sequence collection only (i.e. lncRNA or aTIS and no other database), be identified both by Mascot and Andromeda, to fulfil stringent score cut-offs (see methods), to be backed up by the expression of the underlying transcript in at least one of the tissues (FPKM >1) and to fail a BLAST search against UniProt to exclude obvious alternative explanations. This approach yielded 5 lncRNA and 344 aTIS peptides, respectively. Because of the size of the searched database (aTIS: 474,991 entries; lncRNA: 29,524 entries) there was still ample opportunity to generate false positives. Interestingly, not a single lncRNA peptide could be substantiated by synthetic peptides indicating that lncRNA are rarely if at all translated (Bánfai et al. 2012). To validate the candidate aTIS peptides, we compared spectra of endogenous and synthetic peptide reference standards as described above. Only 66 aTIS peptides (including 8 N-terminally acetylated peptides) covering 53 genes and 57 alternative translation start sites could be confirmed in this way (Table EV6). Manual spectrum interpretation yielded 96 aTIS peptides (overlap of 45 to the SA analysis) mapping to 76 genes and 81 alternative translation start sites. In total, we confirmed 117 aTIS peptides mapping to 89 genes and 99 alternative translation start sites, which include 14 peptides from 12 genes reported in previous studies, for example FXR2, RPA1 and CDV3 (Table EV6) (Kim et al. 2014; Branca et al. 2014). Fifty five of the above aTIS peptides represent 5’ N-terminal extensions of the original gene, 32 peptides represent novel (acetylated) N-termini downstream of the canonical start site, 17 represent frame-shifts potentially leading to an entirely new sequence, 5 peptides likely represent upstream ORFs (uORF) with a stop codon before the canonical start site and 8 peptides with mixed annotation (Fig 4D, left panel). The mirror mass spectra in Fig 4E for the endogenous (top) and synthetic (bottom) peptide (ac)ATTQISKDELDELKEAFAK from the actin binding protein plastin-3 (PLS3) provides an example for the detection of a novel N-terminal sequence. For 36 of the peptides representing aTIS, we identified the exact start site as the peptide was N-terminally acetylated (Fig 4D, right panel). Among these, 18 contained an AUG start codon (Met), 8 contained a GUG start codon (Val), 5 a UUG and 4 a CUG start site (both Leu), and one GCG start site (Ala). This confirms the emerging notion that non-AUG translation initiation events are not as infrequent as previously thought and may represent a mechanism to regulate protein expression (Kearse and Wilusz 2017). This study only identified a relatively small number of aTIS events compared to others (Na et al. 2018) implying that enrichment of N-terminal peptides (Gevaert et al. 2003; Kleifeld et al. 2010) is a more efficient way to detect such events systematically but also pointing out that the previous literature may not be free of a substantial number of mistakes.

An important learning from the present systematic analysis of transcriptomes and proteomes of human tissues is that identifying protein variants or novel coding sequences using proteomics is possible but remains very challenging. There were large discrepancies between the results of the two database search engines used (Mascot and Andromeda) (Fig EV3F-G, Appendix Fig S4 and S5) implying that the underlying scoring schemes are not optimized yet for the detection of variants and novel coding regions. At present, synthetic peptide reference spectra appear to be mandatory for validation and manual spectra comparisons still have a role to play (Lee et al. 2018). Neither approach has been followed systematically in the literature so far and, obviously, they are also not without error but clearly more powerful than purely relying on statistical criteria with largely arbitrary cut-offs alone (Nesvizhskii 2014; Dimitrakopoulos et al. 2016; Lee et al. 2018). It appears that even with the latest proteomic technology, proteogenomics currently offers rather small returns on very significant efforts in data generation, analysis and validation and that large improvements will be required to change this situation substantially in the future. It is possible that our filtering criteria were perhaps too strong so that further variants may be present in the data (see Table EV6, all mirror mass spectra are available in proteomeXchange). However, no convincing false discovery rate estimation has been published yet for spectral angle analysis (let alone for manual data analysis), hence we decided to be conservative. Still, the resource presented in this work should be of considerable value for scientists wishing to develop more sophisticated approaches for proteogenomics in the future and the authors think that there is considerable future potential in the use of synthetic peptide references in conjunction with spectral angle analysis particularly for the many chimeric spectra present in classical data dependent proteomic datasets but more so for the rising use of data independent data acquisition regimes.

## Materials and Methods

### Human tissue specimen

The 29 human tissue samples used for mRNA and protein expression analysis were obtained from the Department of Pathology, Uppsala University Hospital, Uppsala, Sweden as part of the sample collection governed by the Uppsala Biobank (www.uppsalabiobank.uu.se/en/). All tissue samples were collected and handled using standards developed in the Human Protein Atlas (www.proteinatlas.org) and in accordance with Swedish laws and regulations. Tissue samples were anonymised in agreement with approval and advisory reports from the Uppsala Ethical Review Board (References # 2002-577, 2005-338 and 2007-159 (protein) and # 2011-473 (RNA)). The need for informed consent was waived by the ethics committee. The list of all tissues along with corresponding sample preparation and measurement information is provided in Table EV1.

### RNA sequencing

Procedures for RNA extraction from tissues, library preparation, and sequencing have already been described (Uhlén et al. 2015). Briefly, pieces of frozen human tissue were embedded in Optimal Cutting Temperature (OCT) compound and stored at −80°C. Cryosections were cut and stained with hematoxylin-eosin for microscopical confirmation of tissue quality and proper representativity. 5-10 cryosections (10 μm) were transferred to RNAse free tubes for extraction of total RNA using the RNeasy Mini Kit (Qiagen). RNA quality was analyzed with an Agilent 2100 Bioanalyzer system with the RNA 6000 Nano LabChip Kit (Agilent Biotechnologies). Only samples of high-quality RNA (RNA Integrity Number ≥7.5) were used for mRNA sample preparation and sequencing. The mRNA strands were fragmented using Fragmentation Buffer (Illumina) and the templates were used to construct cDNA libraries using a TruSeq RNA Sample Prep Kit (Illumina). Gene expression was assessed by deep sequencing of cDNA on Illumina HiSeq HiSeq 2000/2500 system (Illumina) for paired-end reads with a read length of 2 × 100 bases. RNA sequencing data was aligned against the human reference genome (GRCh38, v83) using Tophat2.0.8b. FPKM (fragments per kilobase of exon model per million mapped reads) values were calculated using Cufflinks v2.1.1 as a proxy for transcript expression level. The FPKM values of each gene were summed up in an individual sample and median normalization was applied to evaluate genes expression levels between tissues. A cutoff value of 1 FPKM was used as a lower limit for detection across all tissues.

### Sample preparation and off-line hydrophilic strong anion chromatography (hSAX)

Fresh frozen human tissue samples (parallel cryosections cut simultaneously as those used for RNA extraction, described above) were prepared for LC-MS/MS as described previously (Ruprecht et al. 2017). Briefly, tissue slices were homogenized in lysis buffer (50 mM Tris/HCl, pH 7.6, 8 M urea, 10 mM tris(2-carboxyethyl)phosphin hydrochloride, 40 mM chloroacetamide, protease and phosphatase inhibitors) by bead milling (Precellys 24, Bertin Instruments, France; 5,500 rpm,2 × 20 s, 10 s pause). Protein content was determined using the Bradford method (Coomassie (Bradford) Protein Assay Kit, Thermo Scientific) and 300 μg of the protein extract were used for in-solution digestion with trypsin. For this, the sample was diluted with 50 mM Tris/HCl to a final urea concentration of 1.6 M, and trypsin was added at a 50:1 (w/w) protein to protease ratio. After 4 hours of digestion at 37°C, another aliquot of trypsin was added to reach a final 25:1 (w/w) protein to protease ratio and the sample was incubated overnight at 37°C. In addition, the tonsil sample was subjected digestion using LysC, ArgC, GluC, AspN, LysN and Chymotrypsin (LysC were from Wako, Japan; the other proteases were from Promega, USA). 300 μg of the protein extract prepared as described above were applied in each digestion. The buffers were prepared according to the manufacturer’s protocols. The resulting peptides were desalted and concentrated on C18 StageTips (Rappsilber, Mann, and Ishihama 2007) and fractionated via hSAX off-line chromatography exactly as described previously (Ruprecht et al. 2017). The details of digestion for each tissue are given in the Appendix Table S1.

### Online liquid chromatography tandem mass spectrometry (LC-MS/MS)

Quantitative label-free LC-MS/MS analysis was performed using a Q Exactive Plus mass spectrometer (Thermo Fisher Scientific, Bremen, Germany) coupled on-line to a nanoflow LC system (NanoLC-Ultra 1D+, Eksigent, USA). Peptides were delivered to a trap column (0.1 × 2 cm, packed with 5 μm Reprosil PUR AQ, Dr. Maisch GmbH, Germany) at a flow rate of 5 μl/min for 10 min in 100% solvent A (0.1% formic acid, FA, in HPLC-grade water). After 10 min of loading and washing, peptides were transferred to a 40 cm (75-μm inner diameter) analytical column, packed with 3 μm, ReproSil-Pur C18-AQ, Dr. Maisch GmbH, Germany) and separated using a 110-min gradient from 2% to 32% solvent B (0.1% FA, 5% dimethyl sulfoxide in acetonitrile, ACN) at a flow rate of 300 nL/min. Full scans (m/z 360-1,300) were acquired at a resolution of 70,000 using an AGC target value of 3e6 and a maximum ion injection time of 100 ms. Internal calibration was performed using the signal of a DMSO cluster as lock mass (Hahne et al. 2013). Tandem mass spectra were generated for up to 20 precursors by higher energy collisional dissociation (HCD) using a normalized collision energy of 25%. The dynamic exclusion was set to 35 seconds. Fragment ions were detected at a resolution of 17,500 using an AGC target value of 1e5 and a maximum ion injection time of 50 ms.

LysC, ArgC, GluC, AspN, LysN and Chymotrypsin digested samples were measured on a Q Exactive HF mass spectrometer (Thermo Fisher Scientific, Bremen, Germany) coupled on-line to a nanoflow LC system (NanoLC-Ultra 1D+, Eksigent, USA). Full scan MS spectra were acquired at 60,000 resolution and a maximum ion injection time of 25 ms. Tandem mass spectra were generated for up to 15 peptide precursors and fragments detected at a resolution of 15,000. The MS2 AGC target value was set to 2e5 with a maximum ion injection time of 100 ms. The other settings were the same as for the Q Exactive Plus.

Tryptic peptides from the tonsil sample were also analysed on an Orbitrap Fusion Lumos Mass Spectrometer (Thermo Fisher Scientific, Bremen, Germany) coupled on-line to a nanoflow LC system (UltiMate™ 3000 RSLC nano System, Thermo Fisher Scientific) using CID, and EThcD/ETD fragmentation. Full MS scans were performed at a resolution of 60,000, a maximum injection time of 50 ms and an AGC target value is 5e5, followed by MS2 events with a duty cycle of 2s for the most intense precursors and a dynamic exclusion set to 60 seconds. CID scans were acquired with 35% normalized collision energy and Orbitrap readout (1e5 AGC target, 0.25 activation Q, 20 ms maximum injection time, inject ions for all available parallelizable time enabled, 1.3 m/z isolation width). EThcD/ETD scans used charge-dependent parameters: 2+ precursor ions were fragmented by EThcD with 28% normalized collision energy and 3+ to 7+ precursor ions were fragmented by ETD. The MS2 scans were read out in the Orbitrap (1e5 AGC target, 0.25 activation Q, and 100 ms maximum injection time).

### MS Data processing and database searching

For peptide and protein identification and label free quantification, the MaxQuant suite of tools version 1.5.3.30 was used. The spectra were searched against the Ensembl human proteome database (release-83, GRCh38) with carbamidomethyl (C) specified as a fixed modification. Oxidation (M) and Acetylation (Protein N-Term) were considered as variable modifications. Trypsin/P was specified as the proteolytic enzyme with 2 maximum missed cleavages. The match between runs function was enabled, with match time window set to 0.7 min and an alignment time window of 20 min. The FDR was set to 1% at both PSM and protein level. LysC/P, ArgC and LysN were specified with 2 maximum missed cleavages. Searches for GluC and AspN peptides allowed 3 missed cleavages. Chymotrypsin (C terminal of F, Y, L, W, or M) was allowed with at most 4 missed cleavages. Label free quantification was performed using the iBAQ approach (Schwanhäusser et al. 2011). For non-tryptic peptides and single tissue analysis, matching data between fractions was disabled.

### Quantitative analysis of transcriptomes and proteomes

The quantitative analyses of proteomic and transcriptomic data were performed at the gene level. To evaluate gene expression level, the total abundance of each gene in all individual sample was used. The data was log transformed (base 10) and normalized using median centering across tissues. The genes were classified into ‘Tissue enriched’, ‘Group enriched’, ‘Tissue enhanced’, ‘Expressed in all’ and ‘Mixed’ as described by Uhlén et al (Uhlén et al. 2015, 2016). Gene ontology analysis of genes only identified in transcriptomes and proteomes, and the elevated proteins expressed in each tissue was performed using the R package ‘clusterProfiler’ and p-values were adjusted according to the method by Benjamini-Hochberg (BH)(Yu et al. 2012). The resulting (redundant) gene ontology terms (biology process) of elevated genes were removed using the ‘simplify’ function in clusterProfiler based on GOSemSim (Yu et al. 2010). The list of 1,158 mitochondrial genes was obtained from MitoCarta 2.0 (Calvo, Clauser, and Mootha 2016). Essential genes (n=583) were assembled from three human essential genes studies using CRISPR-Cas9 and retroviral gene-trap genetic screens (T. Wang et al. 2015; Hart et al. 2015; Blomen et al. 2015). Diseases related genes (n=3,896) and kinase genes (n=504) were obtained from Uniprot. Cancer genes (n=719) were downloaded from Cosmic (Futreal et al. 2004). Drug target genes (n=784) were obtained from Drugbank (Wishart et al. 2018) and restricted to proteins directly related to the mechanism of action for at least one of the associated drugs. GPCR genes (n=1,410) were obtained from HGNC and phosphatase genes (n=238) were from DEPOD (Duan, Li, and Köhn 2015). Transcription factor genes (IF, n=1,639) were obtained from the HumanTFs collection (Lambert et al. 2018).

The Spearman correlation coefficient was used for correlating transcriptome and proteome levels of single tissues. The slopes were estimated by ranged major-axis (RMA) regression, which allows errors in both variables and is symmetric, using the R package ‘lmodel2’ (Csárdi et al. 2015). The protein-mRNA Spearman correlation coefficients of 9,485 genes which were at least expressed in 10 tissues at both mRNA and protein level were calculated. Based on the correlation coefficients, KEGG pathway enrichment analysis was conducted using the Kolmogorov-Smirnov test using the R package ‘fgsea’. The p-value of each pathway was adjusted by the Benjamini-Hochberg method and the cutoff significance was set to 0.05. The Co-inertia analysis (CIA) was performed using the ‘cia’ function in the ‘made4’ R-package (Culhane et al. 2005). 9,485 genes which were expressed in at least 10 tissues at both mRNA and protein level were considered and the remaining missing values were replaced with a positive value 1×10^4^ times smaller than the lowest expression value in each dataset.

### Construction of sample-specific protein sequence databases from RNA-Seq data

RNA sequencing data was aligned to the human reference genome (GRCh38, v83) using Tophat2.0.8b. FPKM values were calculated using Cufflinks v2.1.1 as a proxy for transcript expression level. Rvboost was used for variant calling. All transcripts with FPKM>1 were translated into protein sequences and included in the search database. Each tissue was searched against its matched RNA-Seq database using MaxQuant as described above. The match between runs function was disabled. The MaxQuant output data were used for the isoform analysis.

### Exome sequencing and variant calling

The exome of tonsil tissue was enriched using the Agilent SureSelectXT kit (v5) and sequenced on an Illumina HiSeq4000 sequencer. The raw data was aligned to the human reference genome (hg38) using bwa (v0.7.12) and duplicate reads were marked using Picard Tools (v2.4.1). Genomic variants were called and filtered using the GATK (v.3.6) HaplotypeCaller and VariantFitration modules, respectively, according to the best practice guide (https://software.broadinstitute.org/gatk/best-practices/). Furthermore, variants at sites with a read depth <10x were removed. We also removed any I/L variation as these cannot be distinguished by mass spectrometry. The resulting variants were annotated using the Ensembl Variant Effect Predictor (v85). The RNA sequencing data was aligned to the human reference genome (hg38) using STAR aligner (v2.5.2) and duplicate reads were marked using Picard Tools (v2.4.1). Variants were called using the GATK (v.3.6) HaplotypeCaller module, according to the aforementioned best practice guide.

A variant fasta formatted database was created by the ‘customProDB’ package from the exomic variants (X. Wang and Zhang 2013). Mascot searching of the ultra-deep mass spectrometry data was performed against this database together with protein databases from UniProt and Ensembl using the following parameters: peptide mass tolerance set at 10 p.p.m., MS/MS tolerance set at 0.05 Da, carbamidomethylation of cysteine defined as fixed modification, oxidation of methionine and acetylation defined as variable modification. Trypsin, LysC, ArgC and LysN digested peptides allowed up to 2 missed cleavages. AspN digested peptides with up to 3 cleavages were considered. GluC (V8-DE in Mascot search engine) and chymotrypsin digested peptides were allowed to have a maximum of 3 missed cleavages. Resulting PSMs were analysed using Percolator (v3.01) and an overall FDR cut-off of 1% was applied.

A custom python script was used to identify PSMs covering variant sites and showing either the variant or the canonical genotype. All initial candidate variant peptides had meet the following criteria: i) Mascot ion scores of at least 25; ii) a Mascot delta score of at least 10; iii) the peptide must only map to the variant database; iv) the peptide must map to a single genomic position only; v) for missense variants, the peptide must either show the variant amino acid or it must be cleaved according to a novel protease cleavage site arising from the variant; vi) for nonsense variants, the peptide must end at the novel C-terminus. For canonical genotypes, the same criteria were applied except: i) at least one protein the peptide maps to must not be from the variant database; ii) for missense variants, the peptide must show the wild-type amino acid; iii) for nonsense variants, the end of the peptide must be after the novel C-terminus (after nonsense variant sites). The resulting candidate peptides were mapped against UniProt using BLAST to exclude other obvious explanations. To further consolidate the variants peptides and to reduce false positives, peptide identification by MaxQuant was performed in parallel. Using the customized exomic variant database with the same parameters used in Ensembl database searches described above. The list of candidate variant peptides for the spectra angle analysis required the identification by both Mascot and MaxQuant.

### Identification of peptides translated from non-coding regions

A database of products from possible alternative translation initiation sites (aTIS) was constructed by searching the 5’ UTR of GENCODE transcripts (v25) for putative alternative start codons and in-silico translating these “novel coding sequences”. This resulted in 474,991 aTIS “proteins” > 6 amino acids. The lncRNAs products databases was generated by three-frame-translating the GENCODE (v25) lncRNA database, resulting in 29,524 sequences. The standard 29 tissue proteomics data sets were supplemented with two tissues for which only proteome data was available (bone marrow, pituitary gland), in total 48 samples (including replicates of some organs) was searched against concatenated sequence collections comprising the aTIS and lncRNA databases, GENCODE (v25), UniProt (downloaded 03. February, 2017) and sample specific RNA-Seq based databases using Mascot to identify peptides from known proteins. The search parameters were the same as described for the exome variant peptide identification. The resulting PSMs were processed using Percolator and an overall FDR cut-off of 1% was applied. A custom python script was used to identify PSMs from putative translated lncRNAs or aTIS the database. Candidate peptides had to meet the following criteria: i) the PSM must map to a single database only, i.e. aTIS or lncRNA but no any other; ii) the Mascot score must be at least 25; iii) the Mascot delta score must be at least 10; iv) the original underlying transcript must be expressed in at least one of the tissues (RNA-Seq FPKM >1). The resulting PSMs were then mapped against UniProt using BLAST to exclude other explanations for the novel peptide (e.g. peptides arising from a novel tryptic cleavage site due to a genomic variant). To consolidate the list of candidate aTIS and lncRNA peptides and to reduce false positives, the raw MS data was also searched by MaxQuant (using the same parameters as described for searches using Ensembl). Only those peptides were allowed to pass to the stage of spectral contrast angle analysis if they were identified by both Mascot and MaxQuant.

### Validation of variant and non-coding peptides by synthetic reference peptides

All peptides which passed the filter criteria for Mascot described above were synthesized at JPT Berlin using Fmoc-based solid-phase synthesis. The details of peptide synthesis, sample preparation and MS measurement were as described (Zolg et al. 2017). Normalized Spectral contrast angle (SA) analysis was performed to compare endogenous and synthetic peptides using in-house Python scripts (Toprak et al. 2014). Candidates passed if i) they showed SA values of ≥0.7 (Pearson of ~0.9), ii) the endogenous peptide had a Mascot score of 50 or higher or iii) manual spectrum inspection substantiated the candidate peptide sequence assignment. In parallel, the tandem MS spectra of all candidate peptides were also inspected manually.

### Data availability

Transcriptome sequencing and quantification data are available at www.ebi.ac.uk/arrayexpress/experiments/E-MTAB-2836/. The raw mass spectrometric data and the MaxQuant result files are available from PRIDE (accession number: PXD010154).

## Acknowledgements

This work was in part funded by the German Excellence Initiative cluster Center for Integrated Protein Analysis Munich (CIPSM). This work was in part supported by the Knut and Alice Wallenberg Foundation. The authors wish to thank pathologists and staff at the Department of Clinical Pathology, Uppsala University Hospital for providing the tissues used in the study, Harald Marx for guidance on the construction of protein sequence databases and Professor Marily Theodoropoulou for providing pituitary samples. DW is grateful for a scholarship from the China Scholarship Council. BE and JG are supported by EU Horizon2020 Collaborative Research Project SOUND. A Fellowship by the Graduate School of Quantitative Biosciences Munich (QBM) supports BE.

## Author contributions

BK, FP, HH, JG, MU conceived and designed the study

FP selected and provided normal human tissue samples

AA, DW, DZ, KS, JZ performed experiments

BE, BH, BK, CM, DW, HH, JG, MF, MW, TH, TS, TW performed data analysis

BE, BK, DW, HH, JG, TW wrote the manuscript

## Competing financial interests

HH and TH are employees of OmicScouts GmbH. HH, MW and BK are co-founders and shareholders of OmicScouts GmbH. MW and BK have no operational role in OmicScouts GmbH. KS is employee of JPT Peptide Technologies.

---

**Figure EV1.**
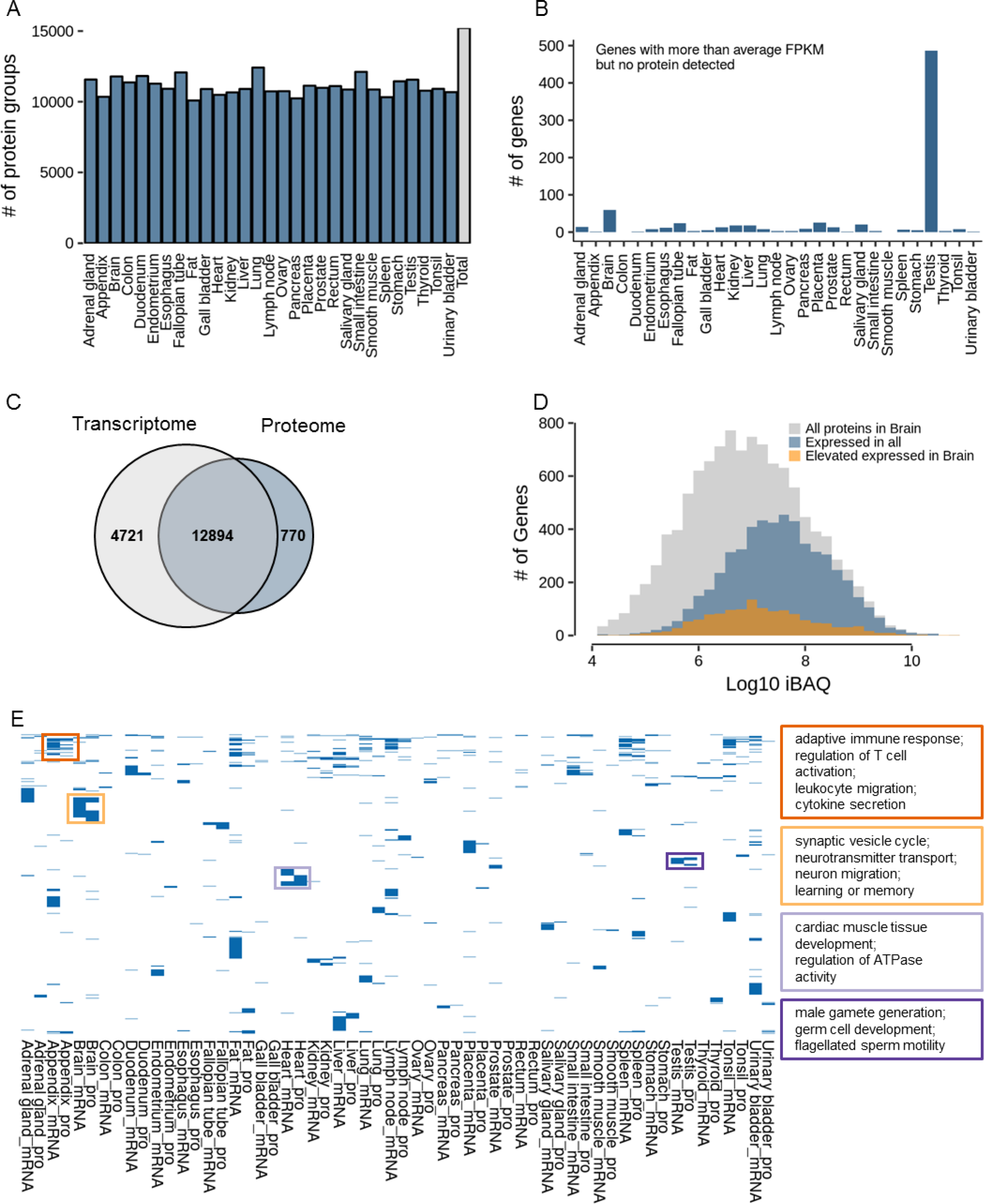
Further characterization of human proteomes and transriptomes. (A) Number of identified protein groups in all 29 tissues. (B) Number of genes in all tissues that were detected at the transcript but not at the protein level. (C) Comparison of genes covered by either transcripts or proteins. (D) Abundance distribution of all proteins detected in human brain (grey). Proteins in blue are expressed in all 29 tissues and proteins in orange show elevated expression in brain. (E) Clustering of gene ontology terms (biological process) for proteins and transcripts that show the most divergent expression across all tissue. Boxes give examples of GO terms for four different tissues (appendix, brain, heart, and testis).

**Figure EV2.**
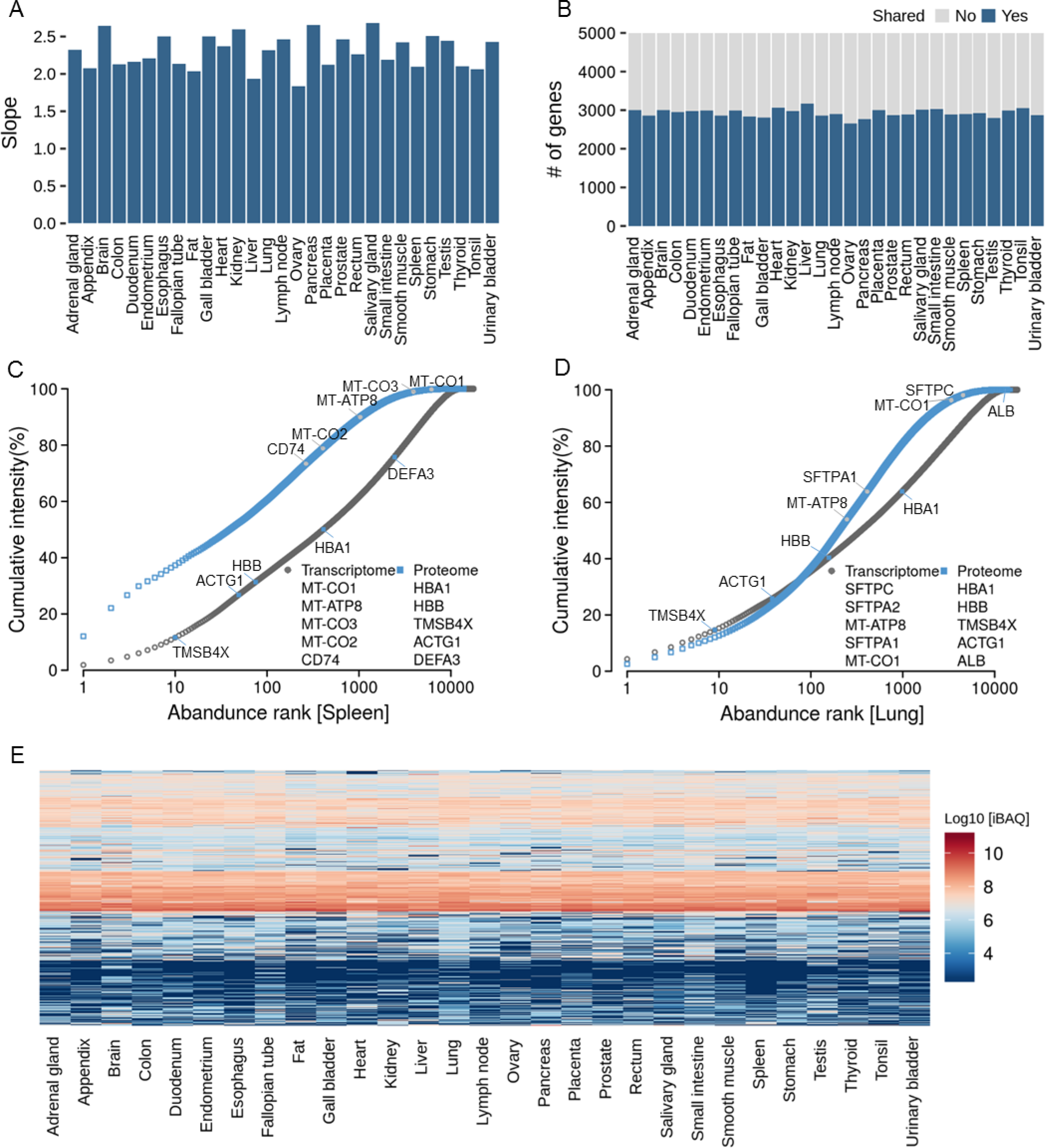
Relationships between mRNA and protein expression. (A) Slopes of protein vs mRNA abundance plots (see main Fig 2B) for all tissues. (B) Number of genes that are shared among the 5,000 most abundant transcripts or proteins in all tissues. (C, D) Ranked abundance plots for transcripts and proteins in spleen and lung showing different characteristics in the abundance distributions (see also main Fig 2C and Appendix Fig S2 for all tissues). (E) Clustering of protein abundances across all tissues. It is apparent that many proteins have similar expression levels across several/many tissues.

**Figure EV3.**
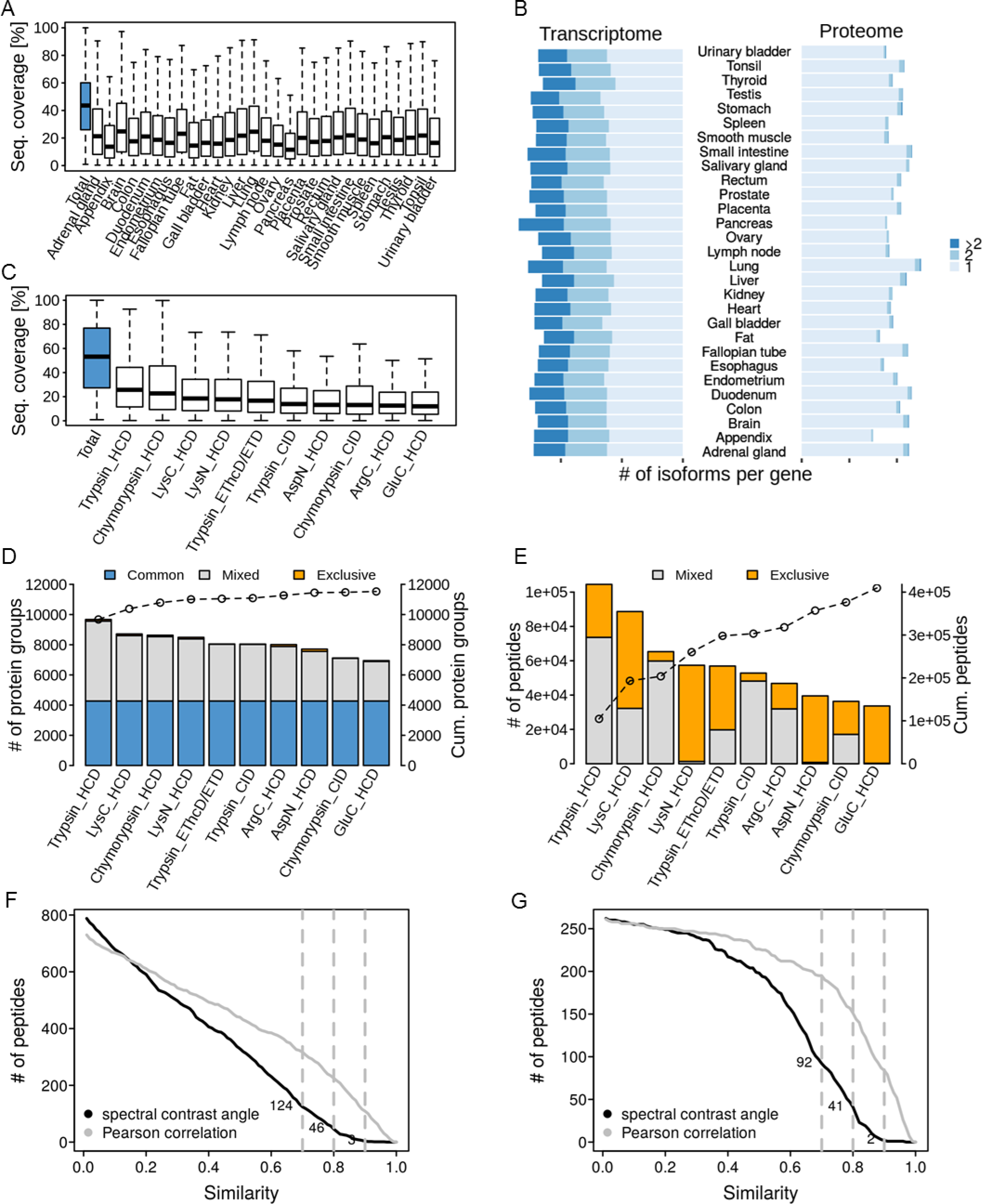
Proteogenomic characterization of human tissues. (A) Distribution of peptide sequence coverage obtained for proteins by mass spectrometry in all tissues. (B) Analysis of the number of isoforms detected by transcriptomics or proteomics in all tissues. (C) Distribution of peptide sequence coverage obtained for proteins by mass spectrometry in tonsil tissue broken down by protease and fragmentation method used. (D) Number of identified proteins broken down by protease and fragmentation method used. Proteins covered by all workflows are marked in blue. The line indicates the cumulative number of proteins when adding data from the individual workflows. (E) Same as panel D but for peptides. Peptides covered by more than one workflow are marked in grey, those exclusive for one workflow are marked in orange. (F) Number of experimental vs synthetic peptide reference spectra comparisons for candidate aTIS peptides (only the spectra with highest spectral angle of each peptide was plotted) after database searching using Mascot as a function of the spectral angle and Pearson correlation coefficient. Dotted lines mark spectral angles of 0.7, 0.8 and 0.9. (G) Same as panel F but showing only candidate peptides that were identified by both Mascot and Andromeda.

**Appendix Table S1.**
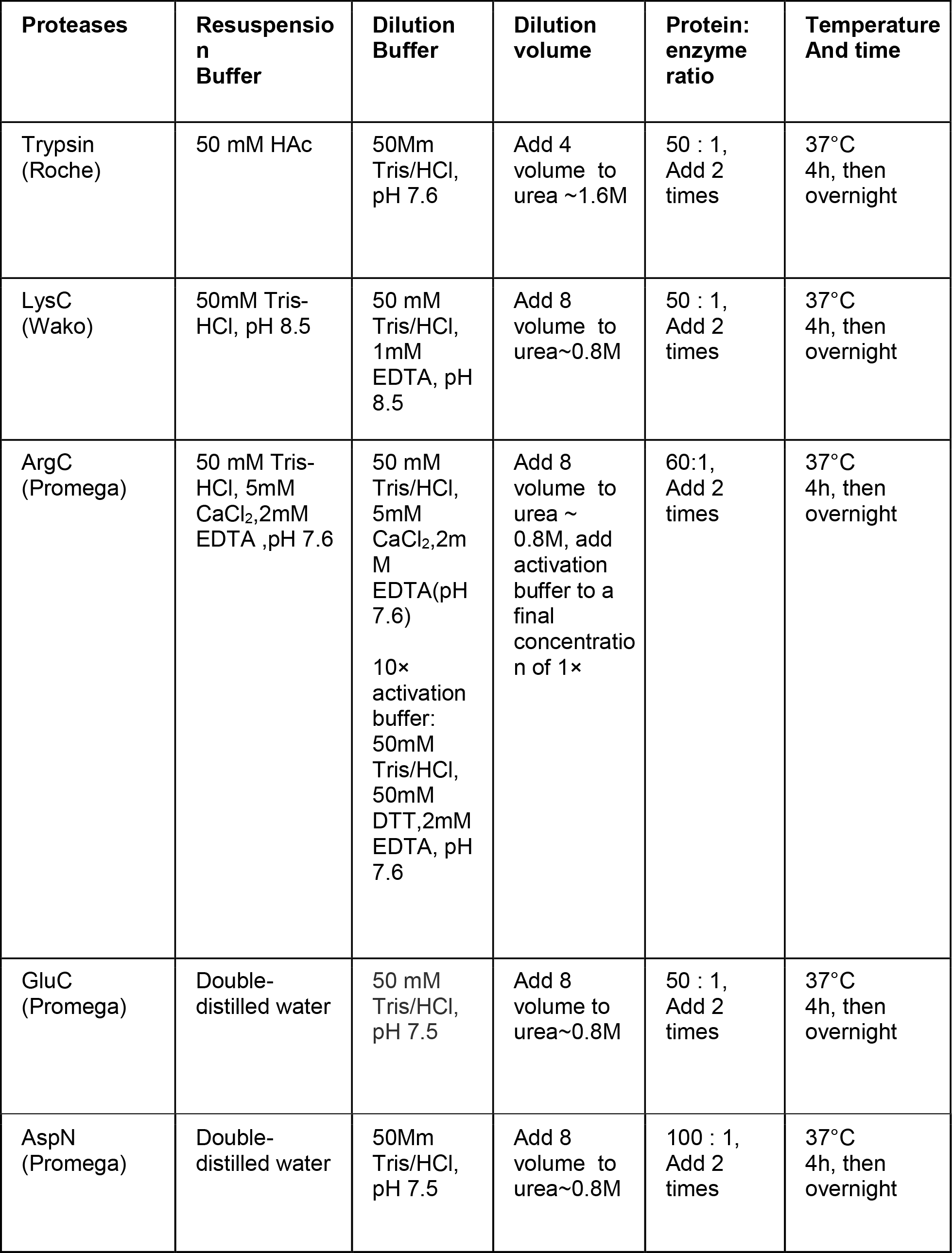
Digestion conditions for each protease

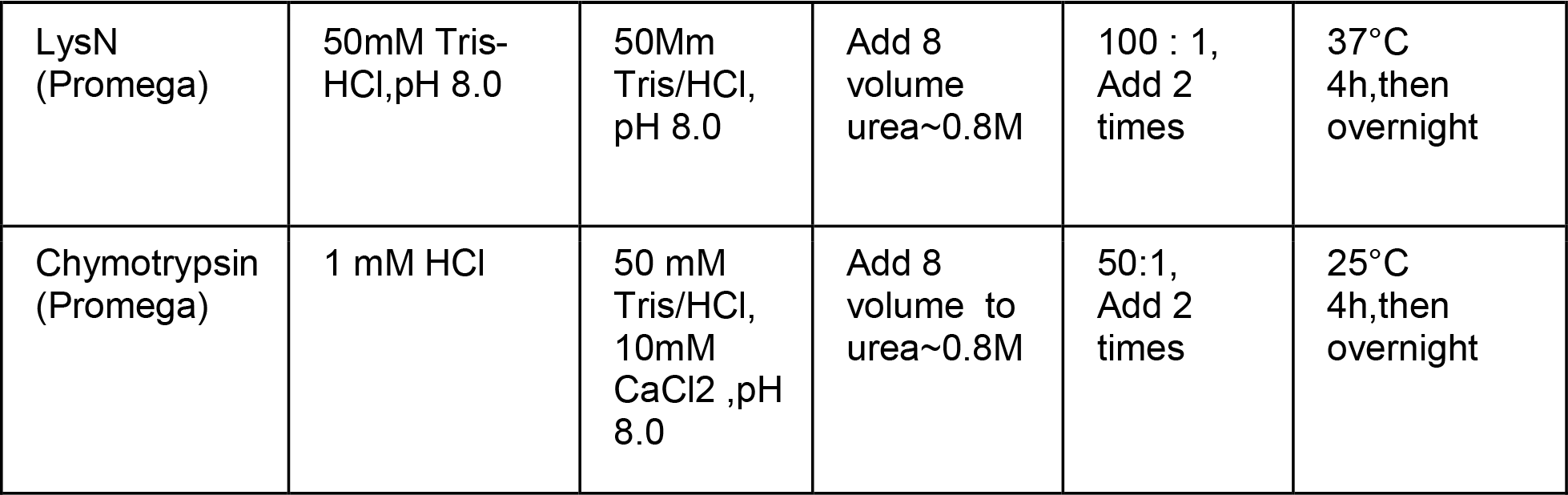

**Appendix Figure S1.**
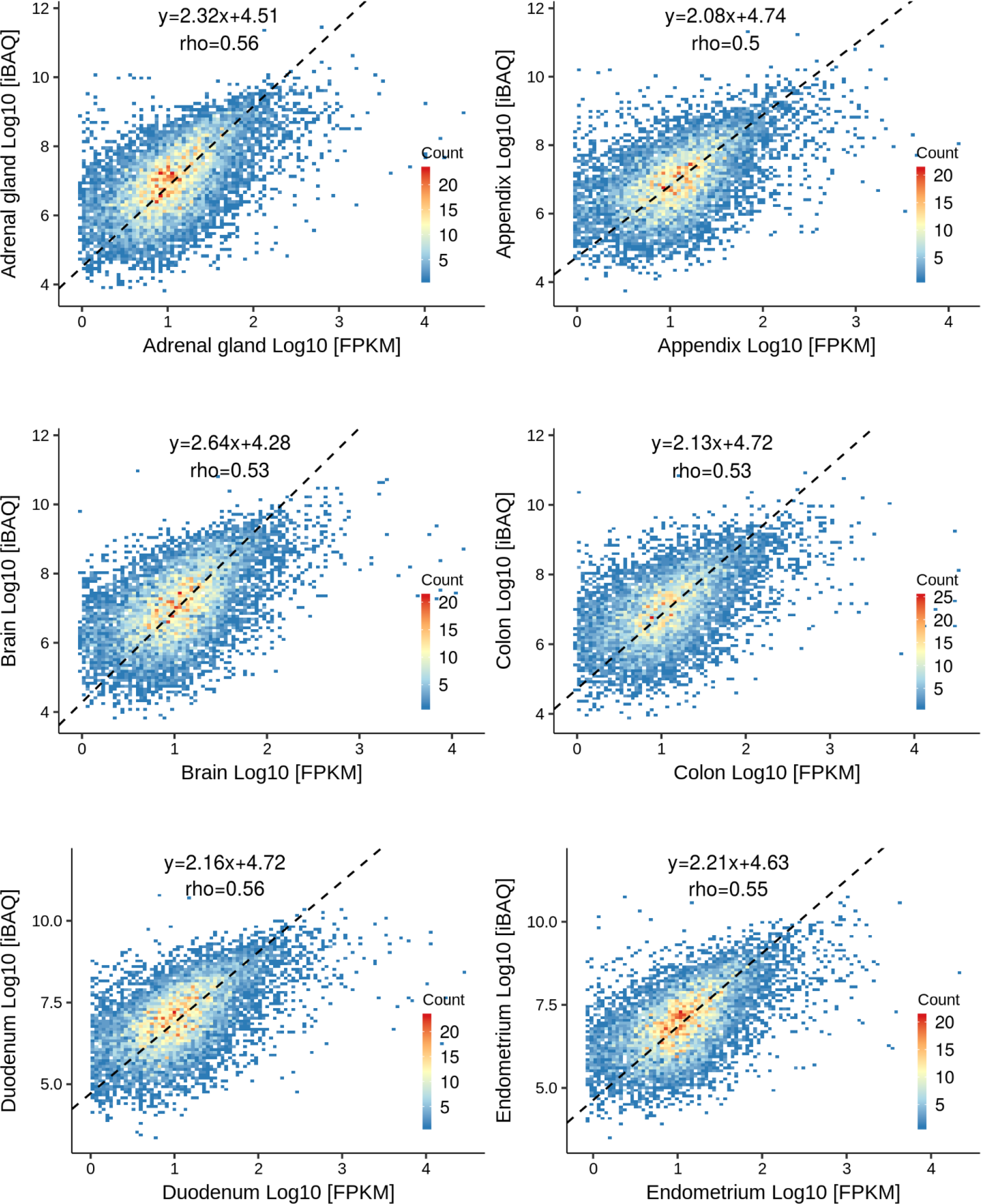
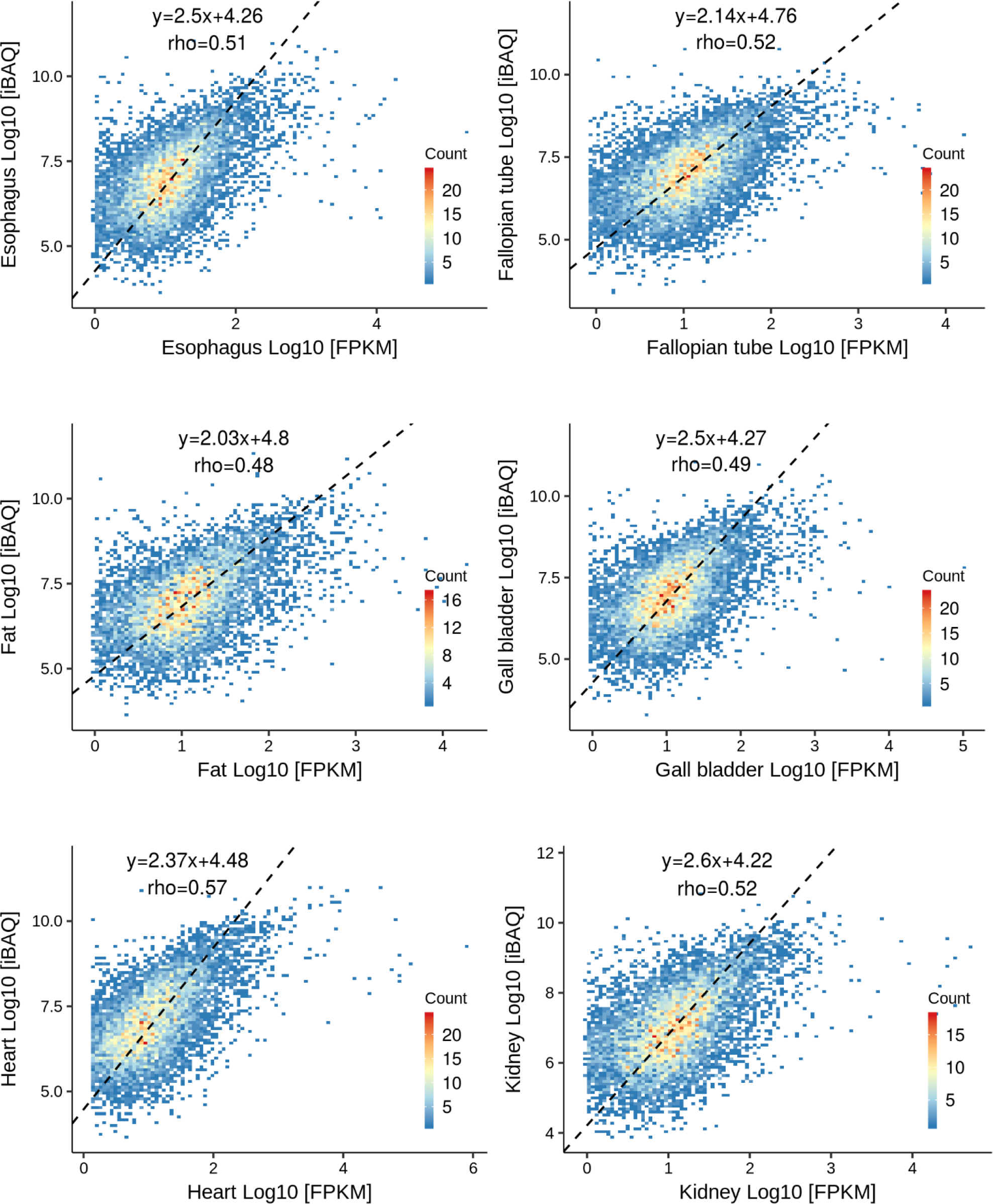
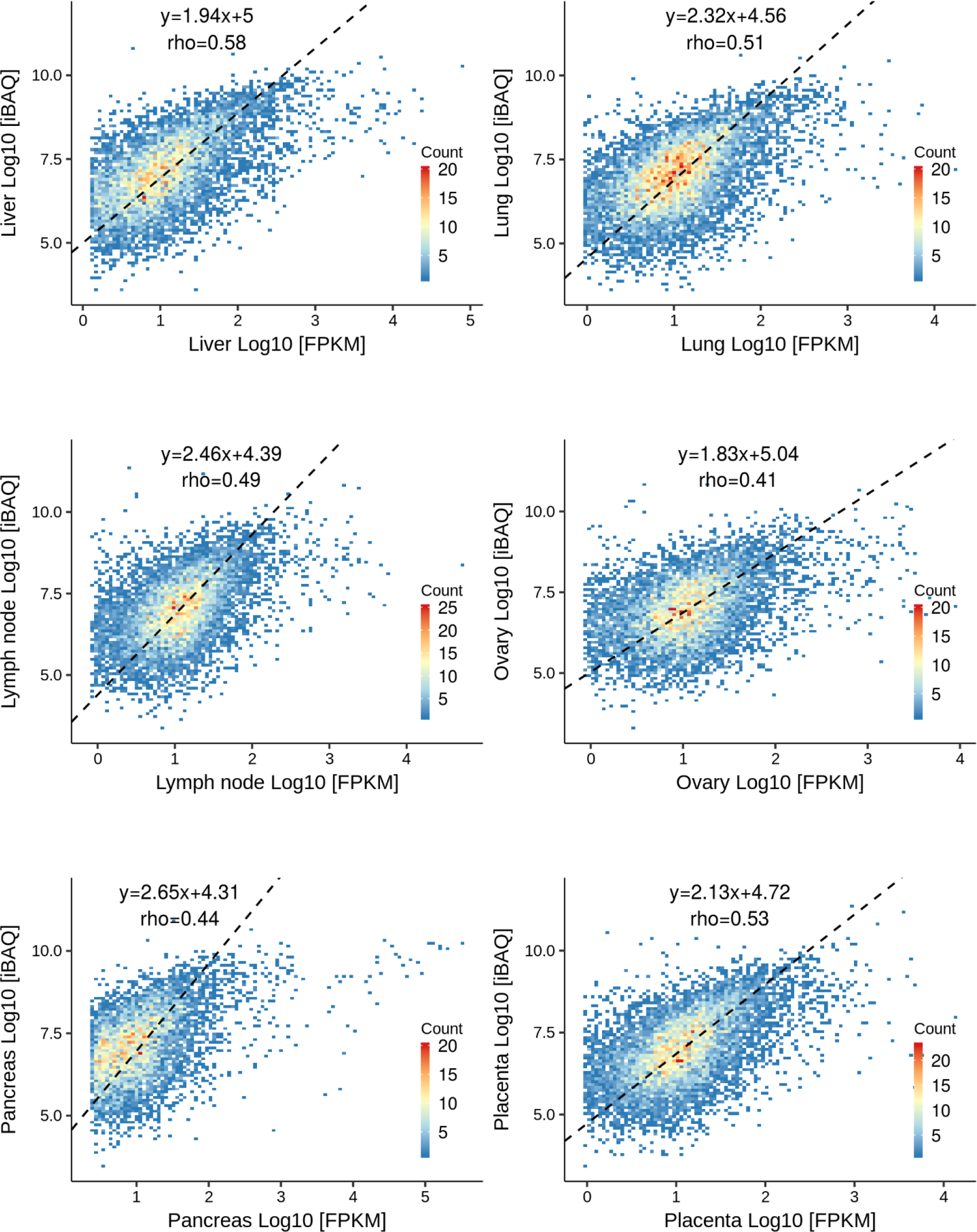
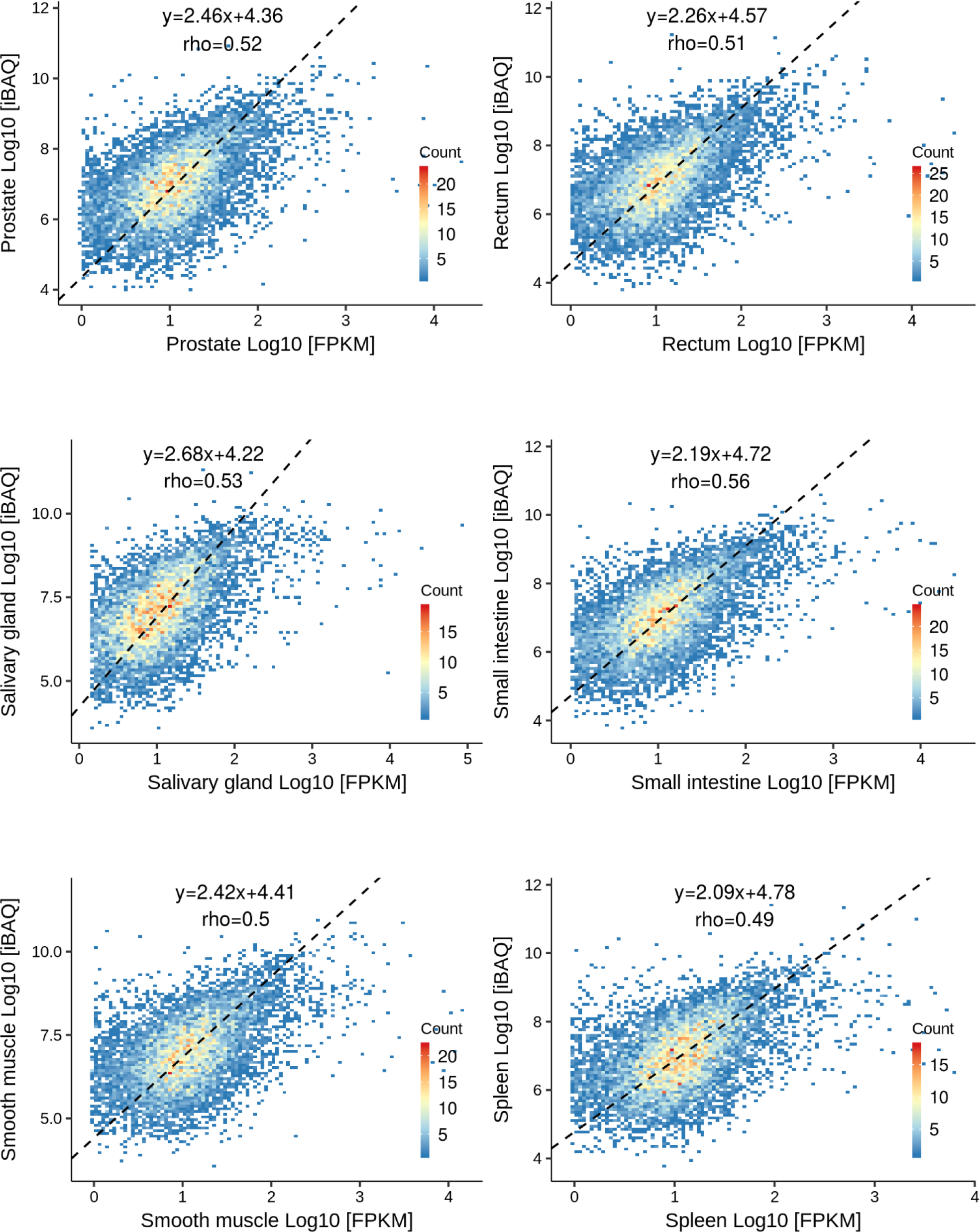
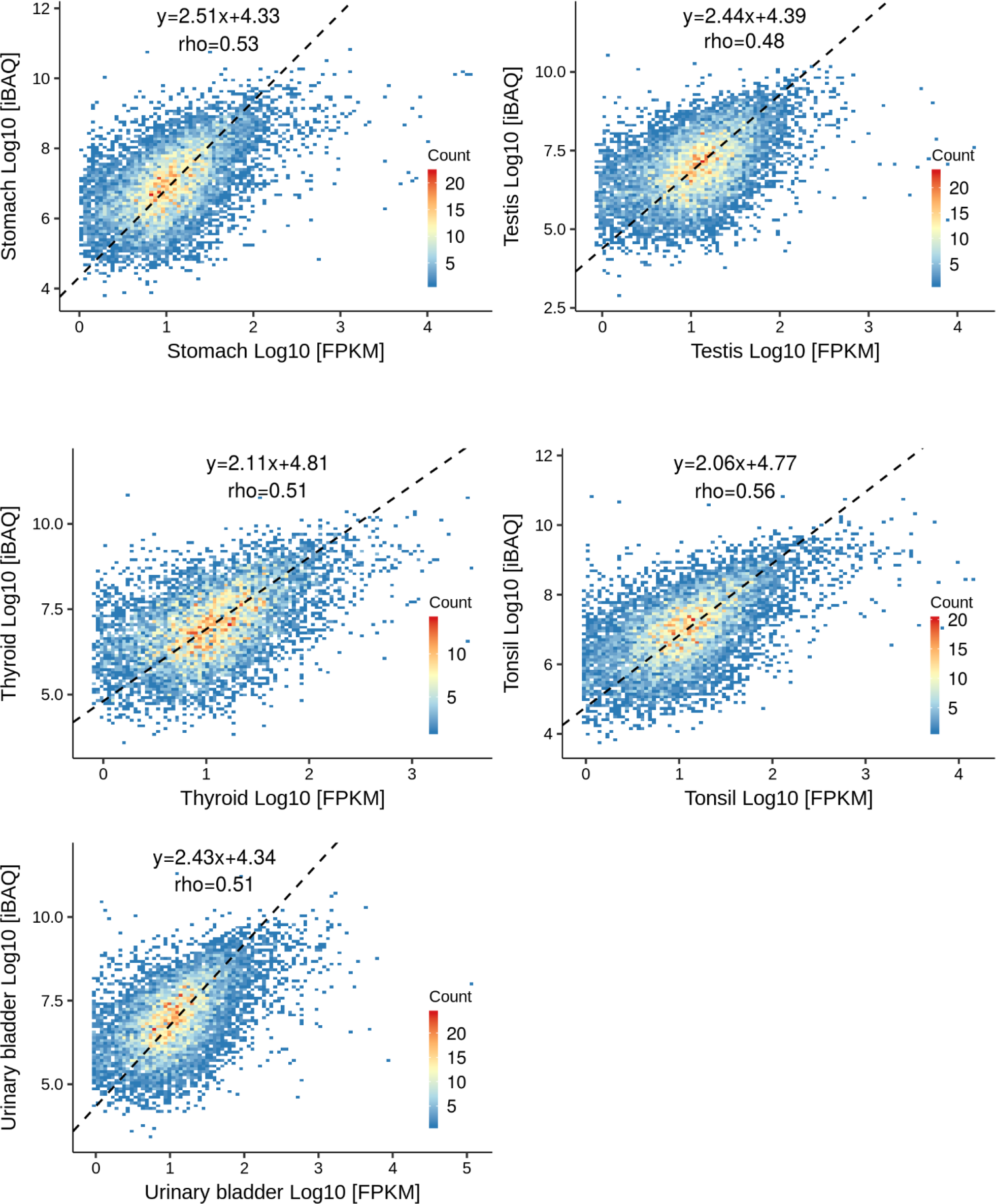
Protein to mRNA abundance plots of 29 tissues

**Appendix Figure S2.**
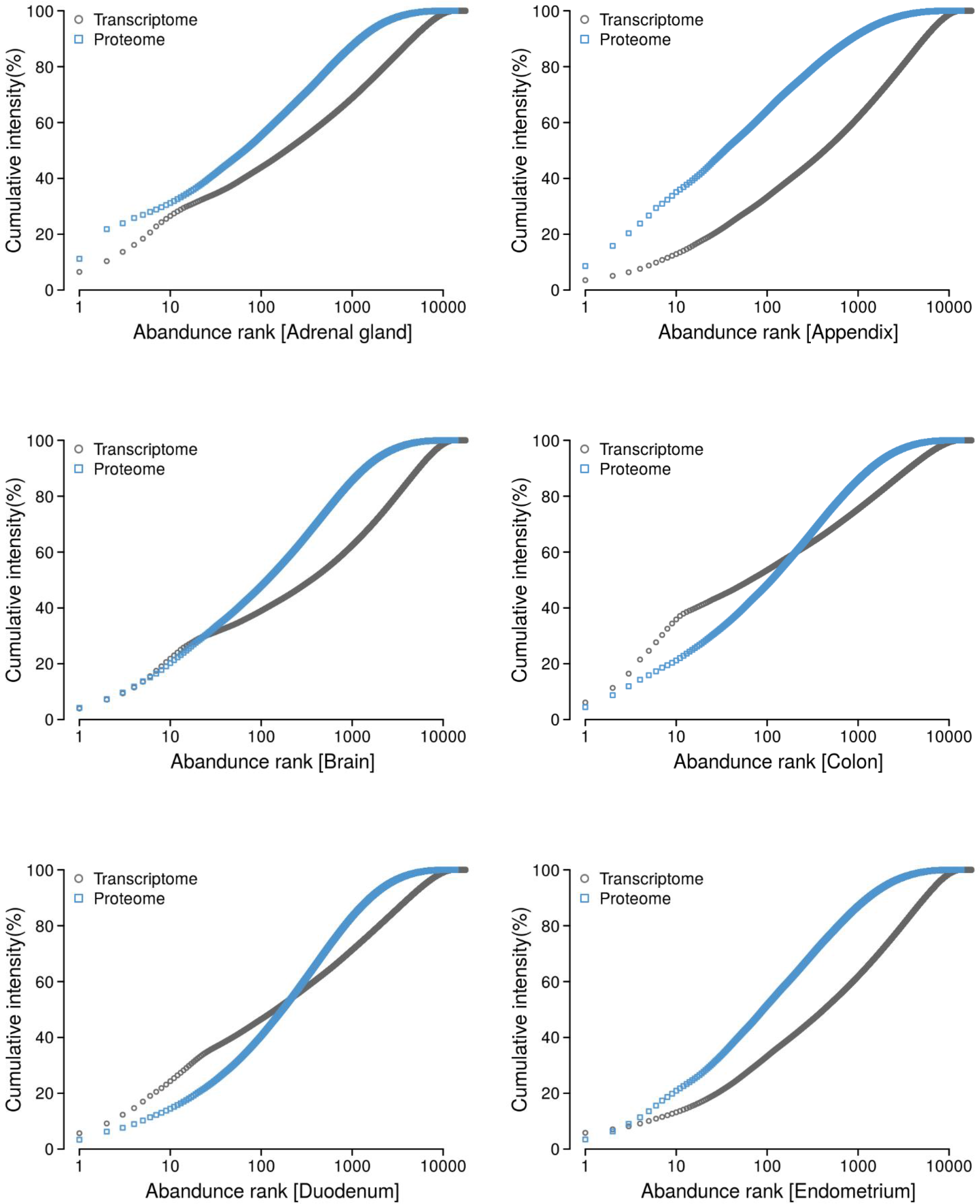
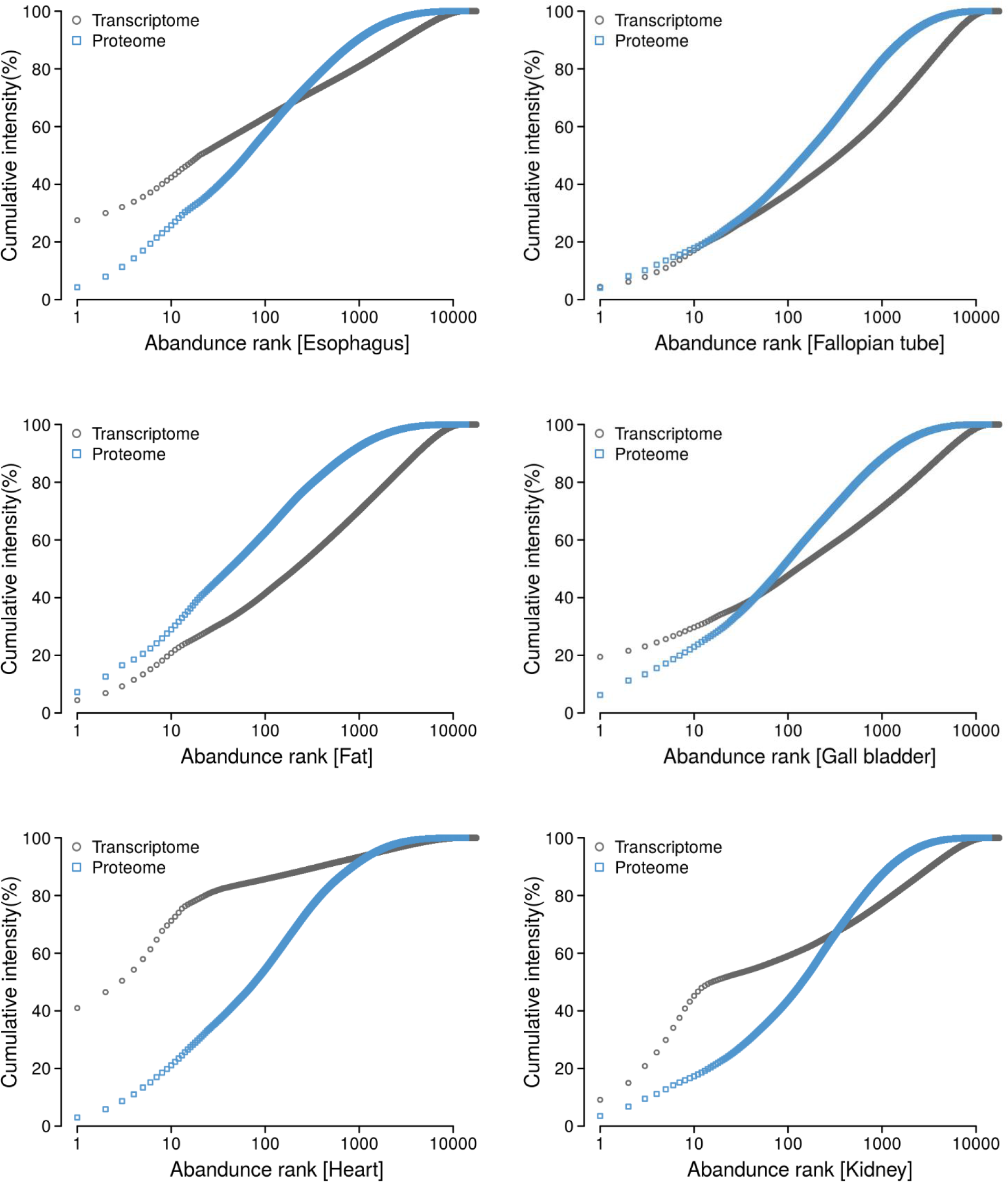
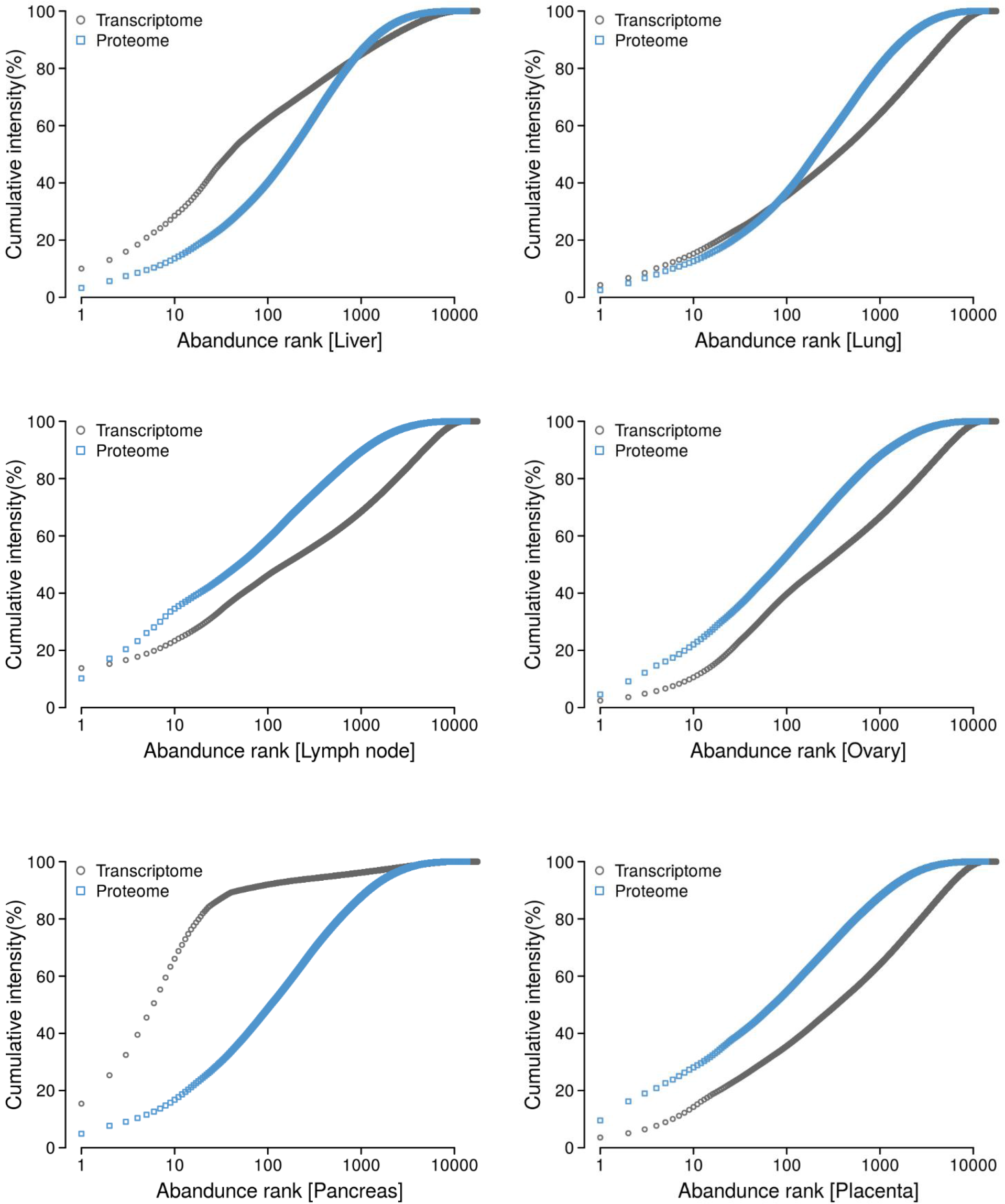
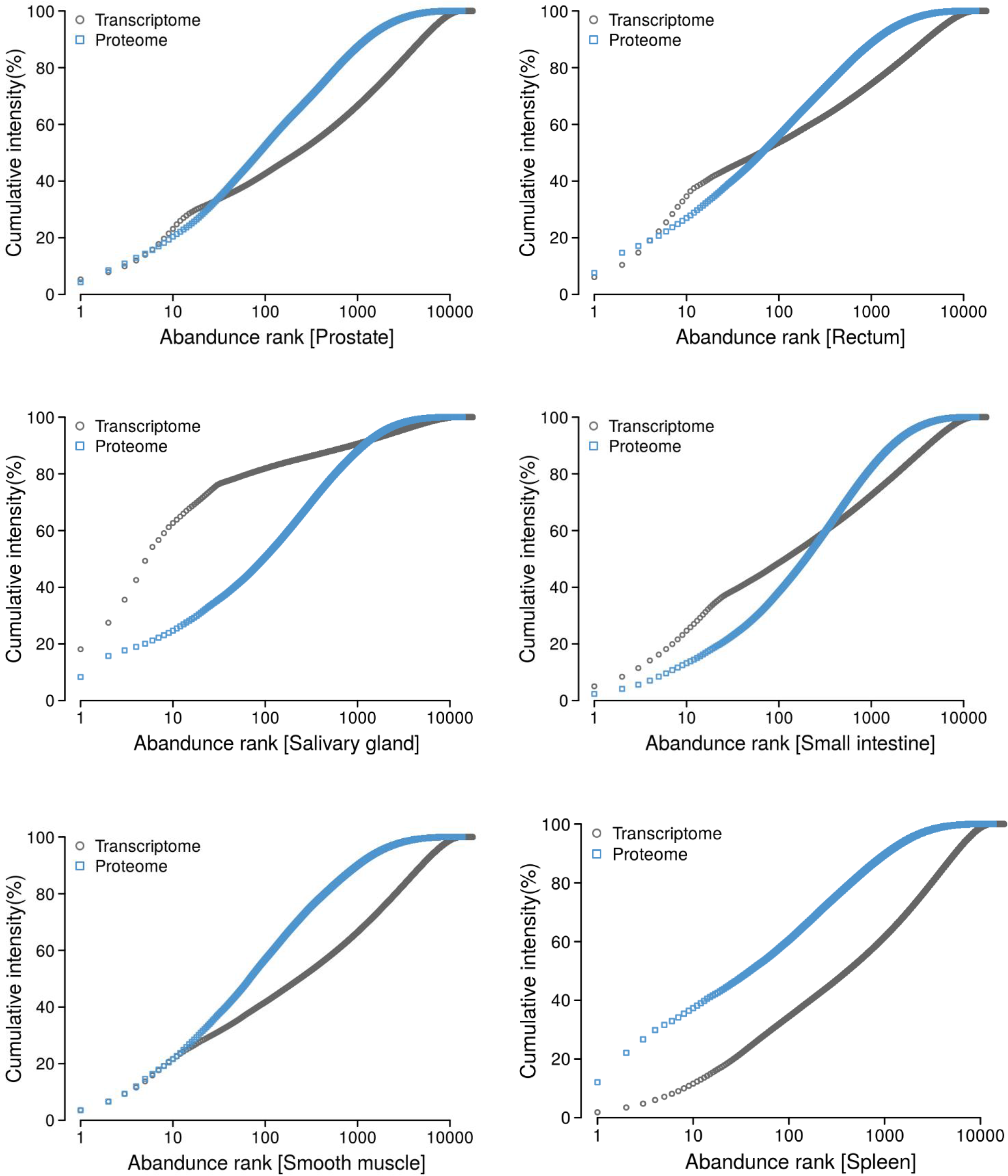
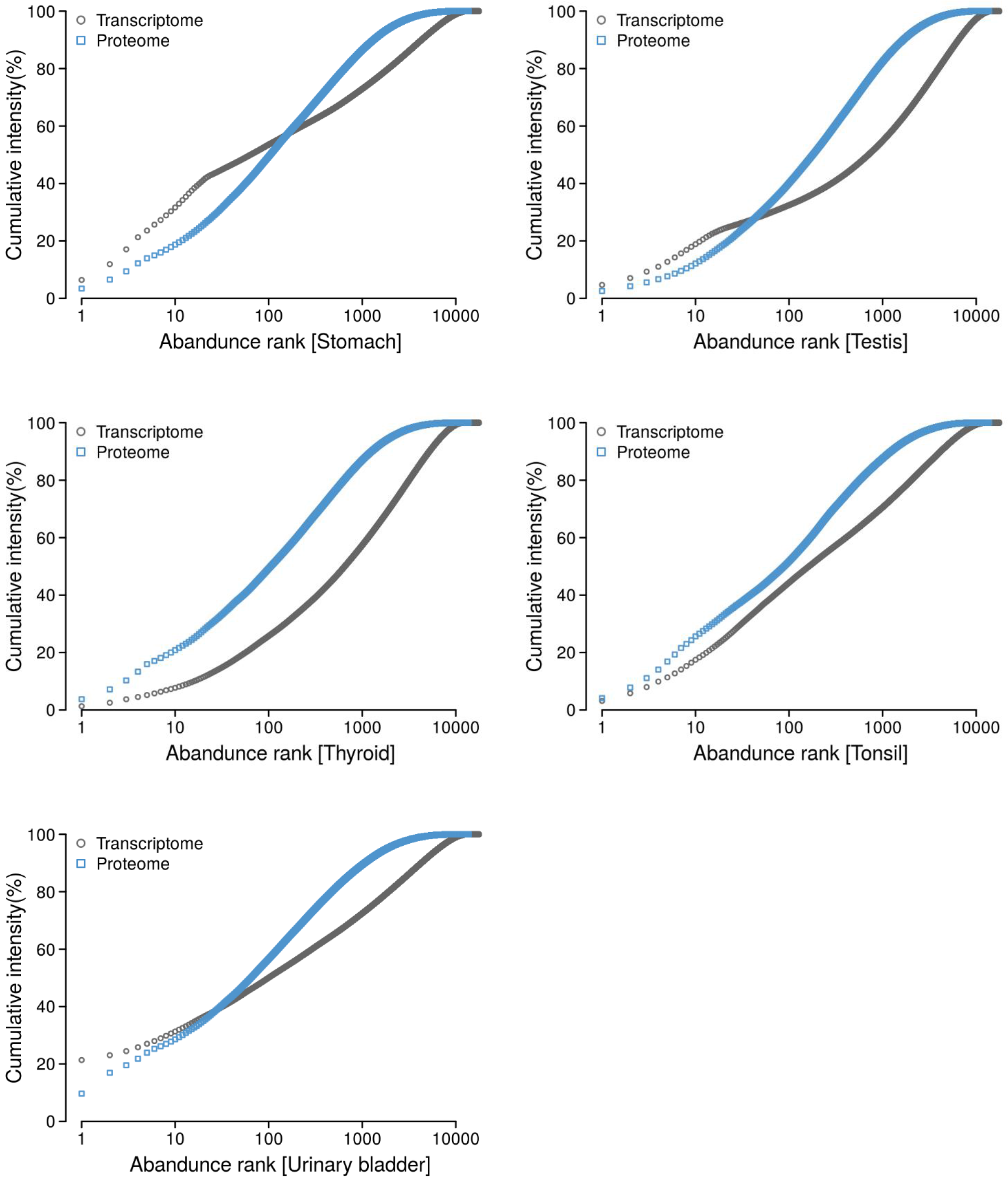
Abundance distribution of all proteins detected in 29 tissues

**Appendix Figure S3.**
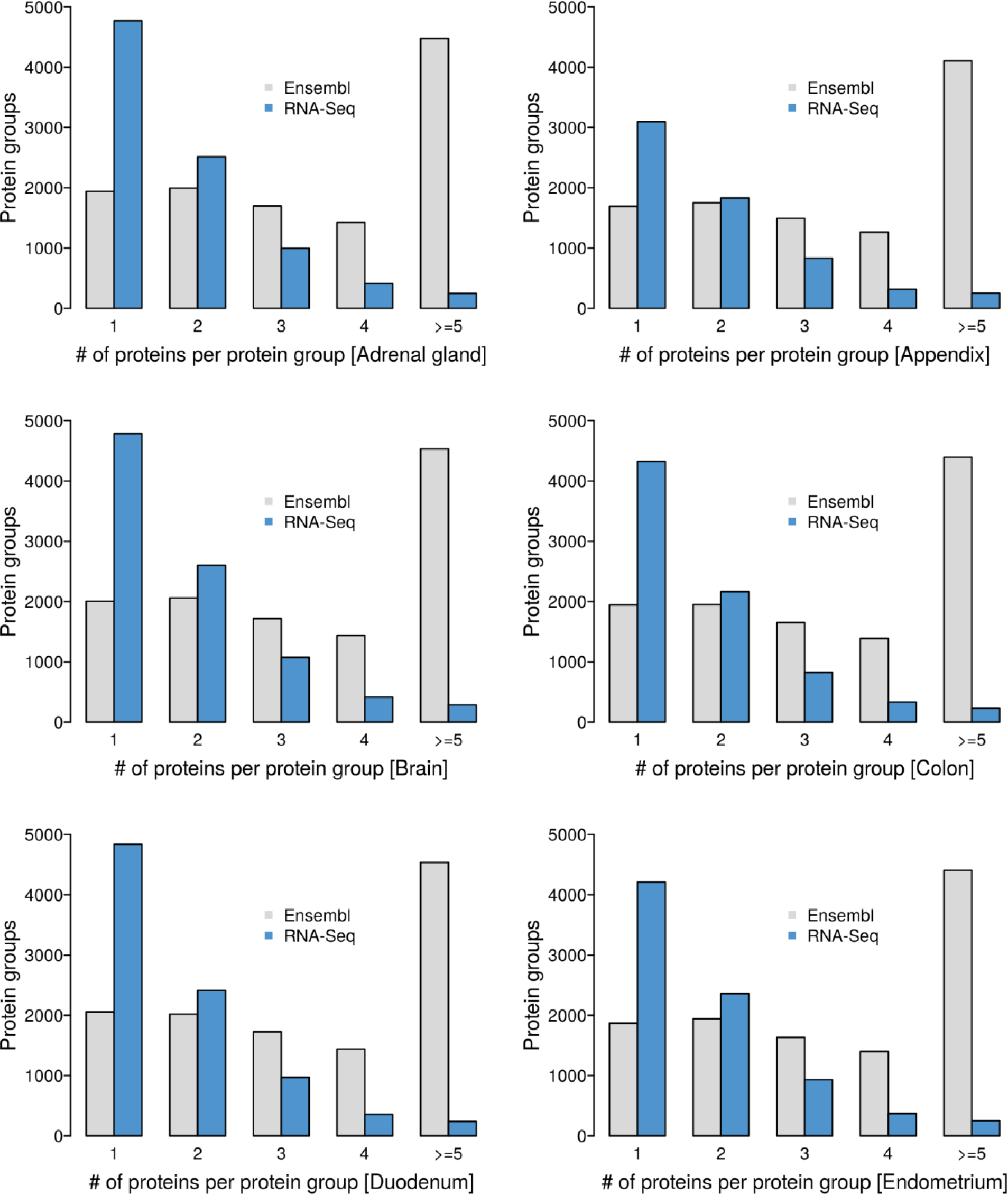
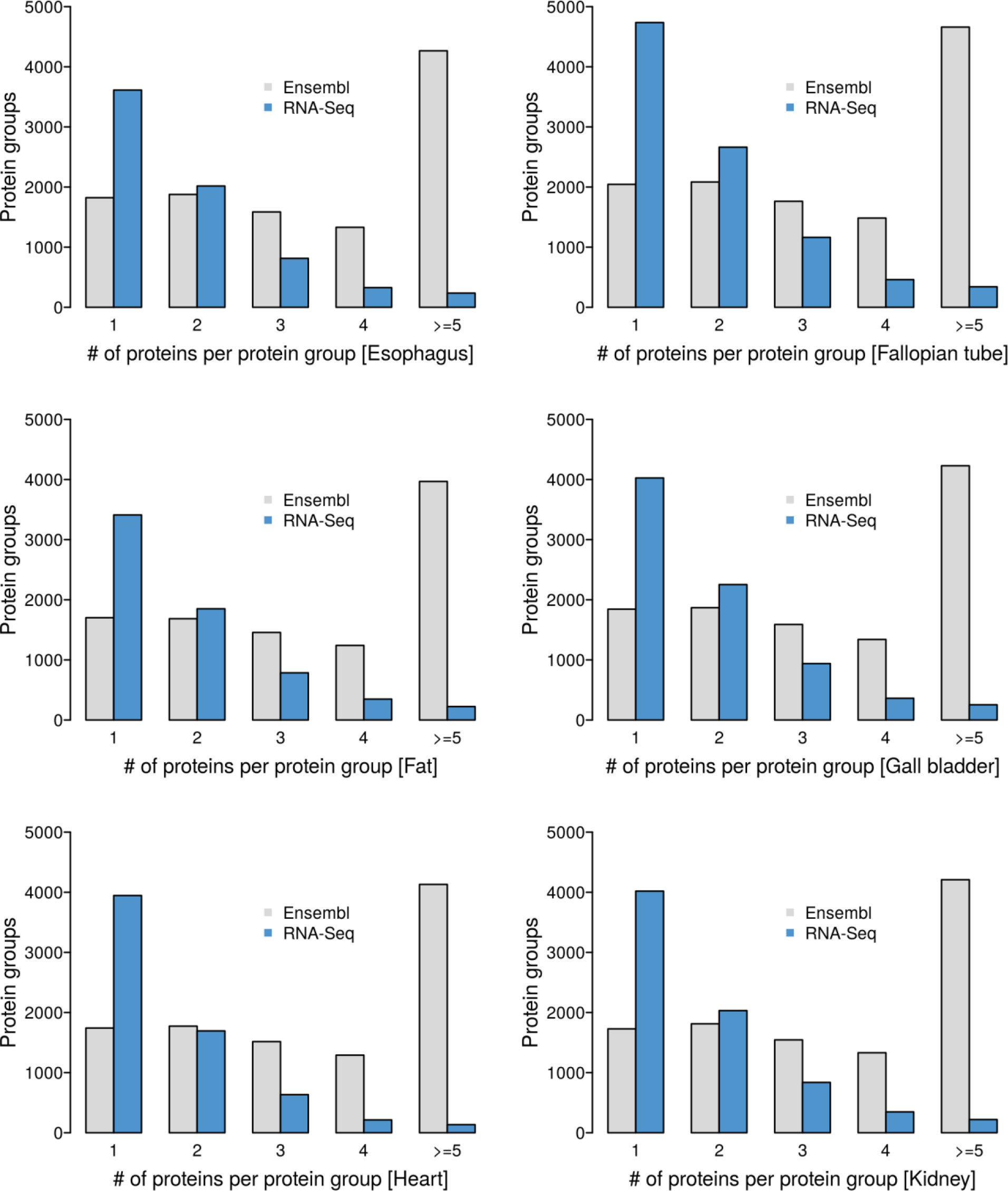
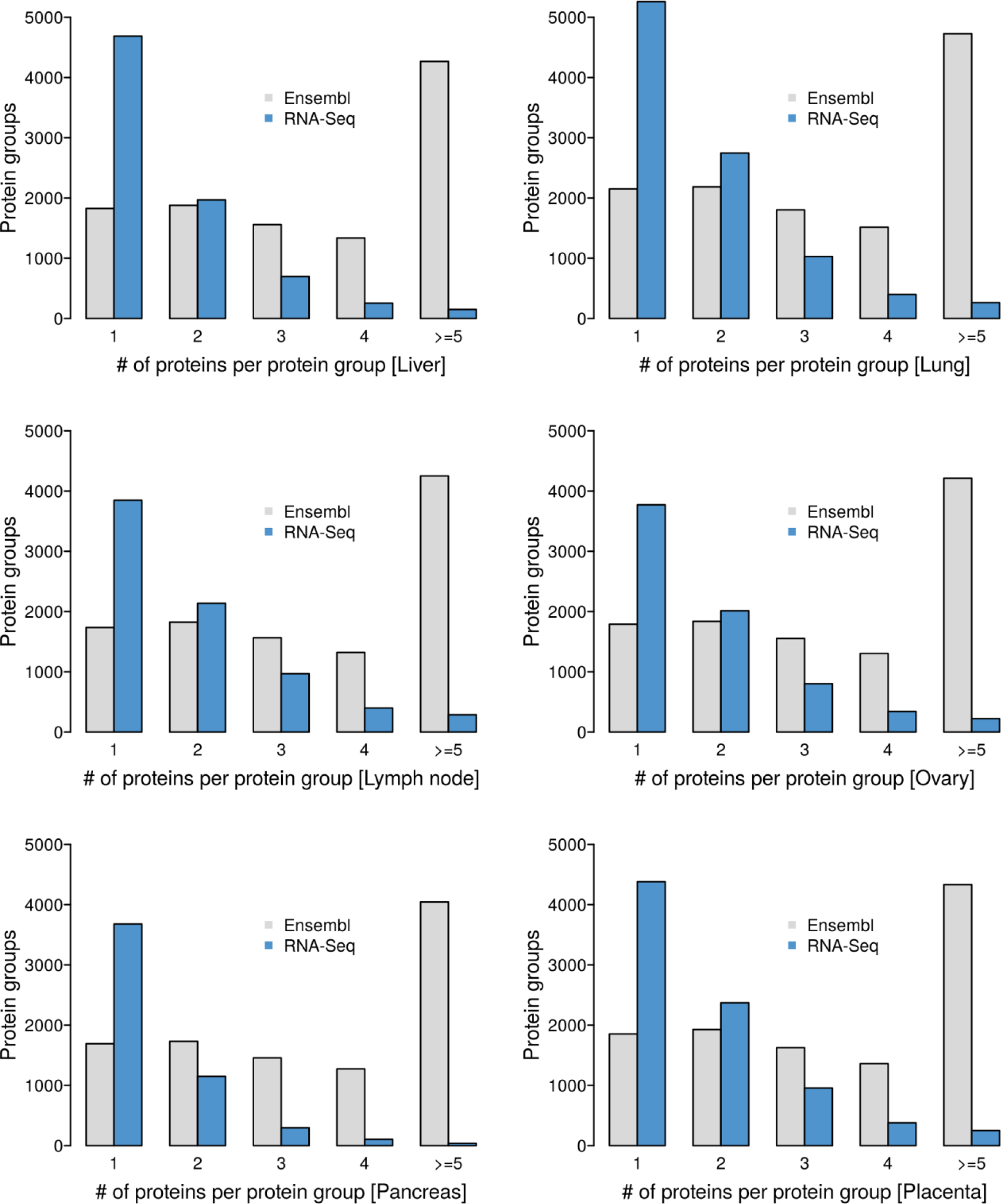
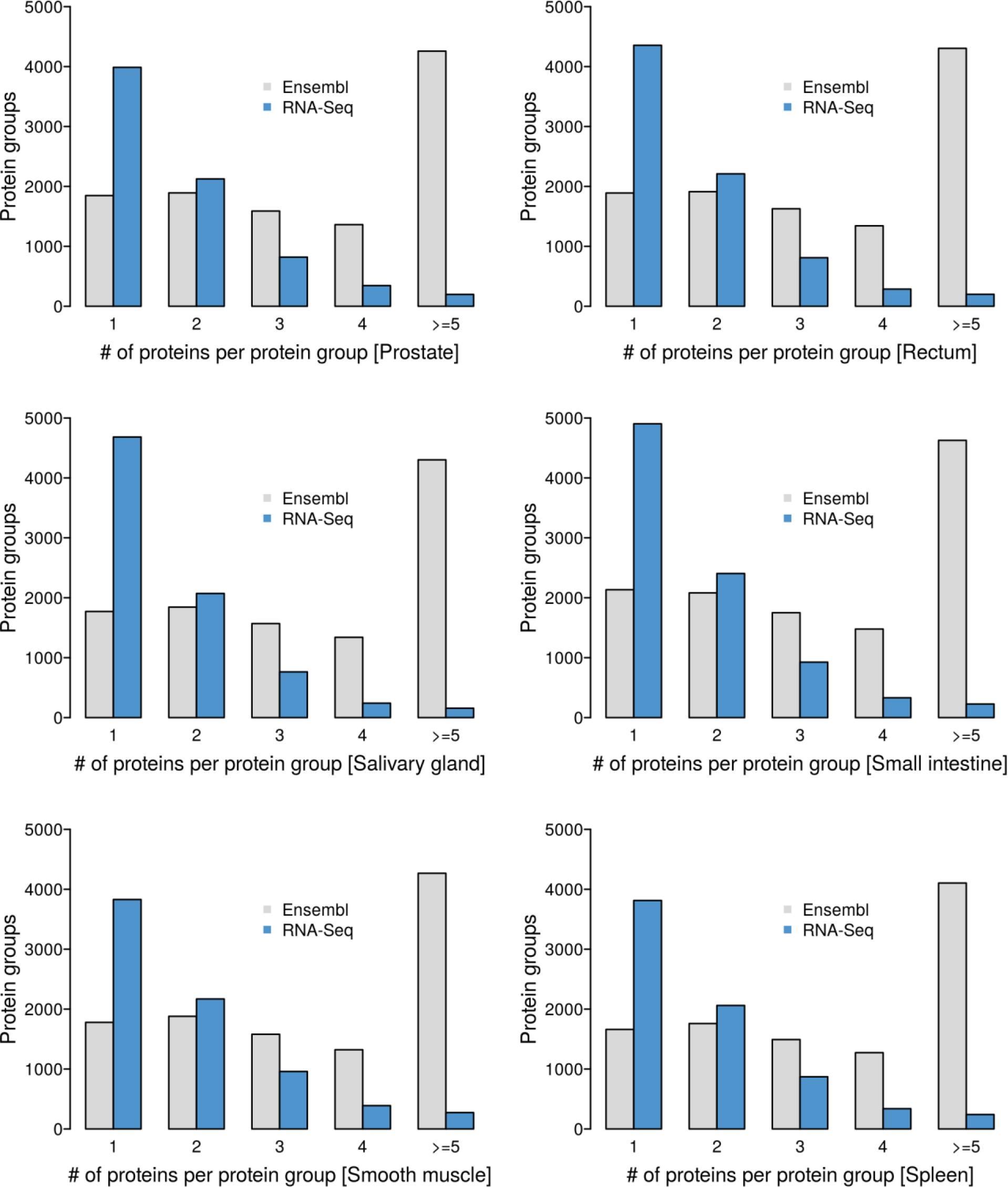
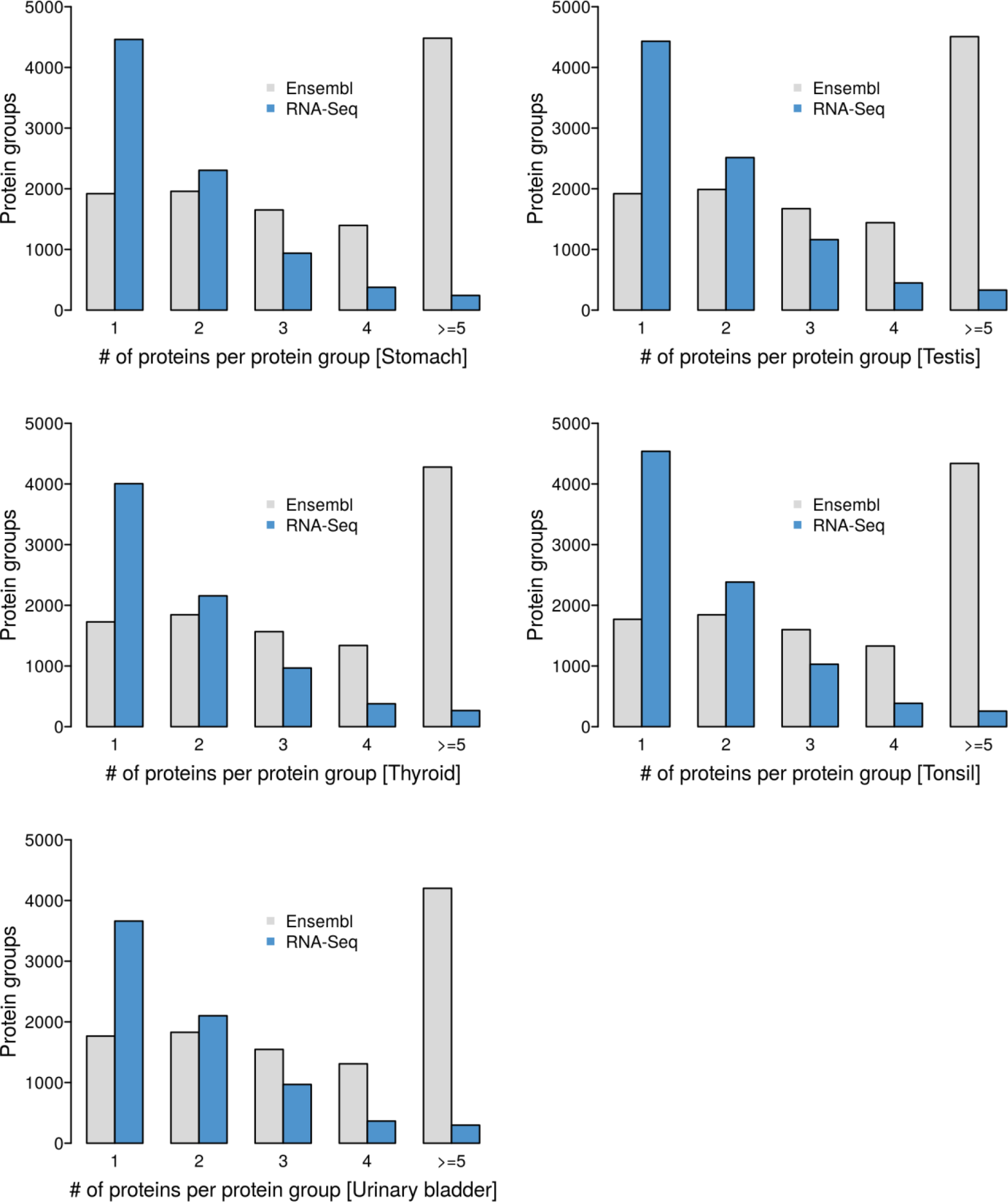
Searching the proteomic data of 29 tissues against their tissue-specific sequence databases constructed from RNA-Seq data drastically reduces the number of individual protein sequences in protein groups compared to searches against Ensembl

**Appendix Figure S4.**
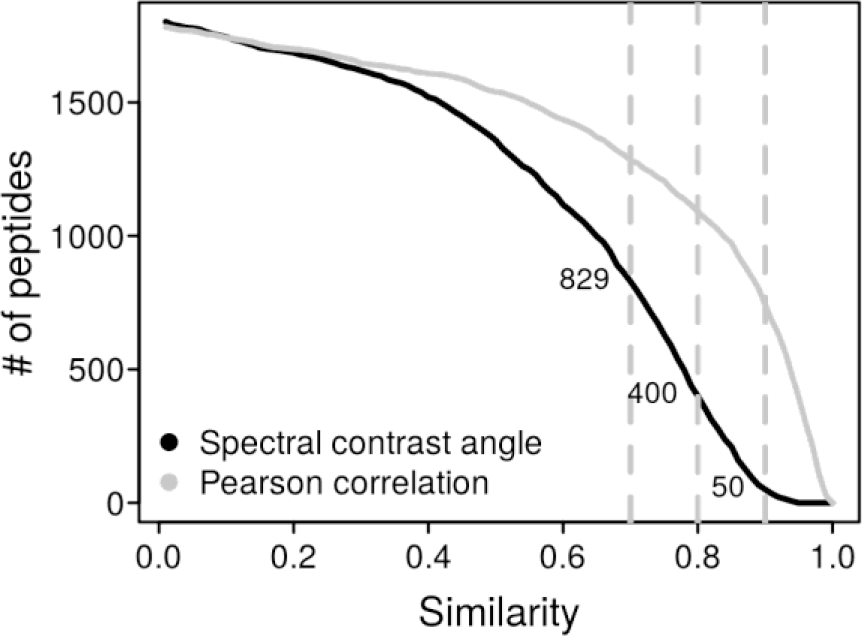
Number of experimental vs synthetic peptide reference spectra comparisons for candidate SAAV peptides (only the spectra with highest spectral angle of each peptide was plotted) after database searching using Mascot as a function of the spectral angle and Pearson correlation coefficient. Dotted lines mark spectral angles of 0.7, 0.8 and 0.9.

**Appendix Figure S5.**
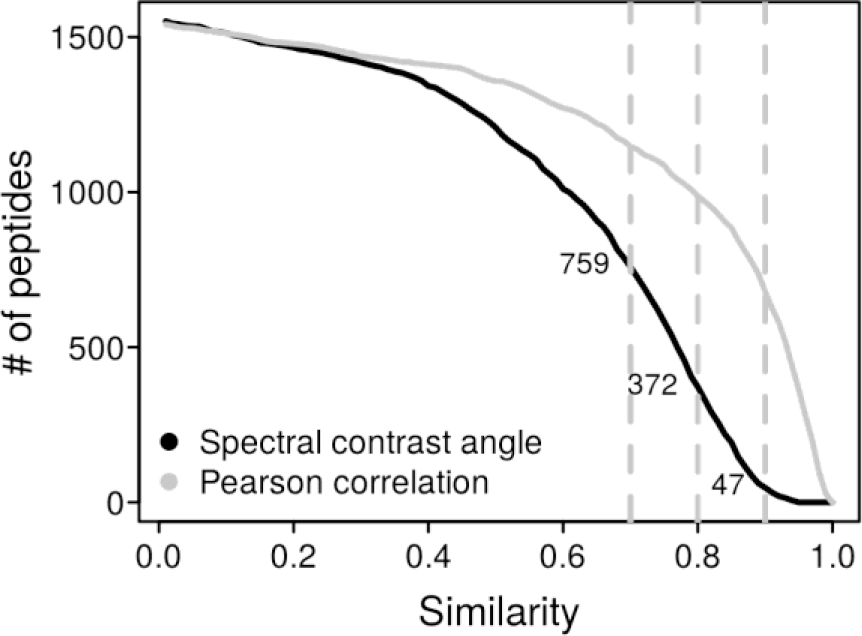
Number of experimental vs synthetic peptide reference spectra comparisons for candidate SAAV peptides identified by both Mascot and Andromeda (only the spectra with highest spectral angle of each peptide was plotted) as a function of the spectral angle and Pearson correlation coefficient. Dotted lines mark spectral contrast angles of 0.7, 0.8 and 0.9.

## Abbreviations

aTIS: Alternative translation initiation sites
BH: Benjamini-Hochberg
CIA: Coinertia analysis
CID: Collision-induced dissociation
ETD: Electron-transfer dissociation
EThcD: Electron-transfer/Higher-energy collision dissociation
FDR: False discovery rate
FPKM: Fragments per kilobase million
GPCR: G-protein-coupled receptors
HCD: Higher-energy collision dissociation
LC-MS/MS: Liquid chromatography tandem mass spectrometry
lncRNA: Long non-coding RNA
MS: Mass spectrometry
PTR: Protein-to-mRNA
SAAV: Single amino acid variant
SNV: Single nucleotide variant
TF: Transcription factor
uORF: Upstream open-reading frame

## References

Bánfai, Balázs, Hui Jia, Jainab Khatun, Emily Wood, Brian Risk, William E. Gundling Jr, Anshul Kundaje, et al. 2012. “Long Noncoding RNAs Are Rarely Translated in Two Human Cell Lines.” Genome Research 22 (9): 1646–57.

Becher, Isabelle, Thilo Werner, Carola Doce, Esther A. Zaal, Ina Tögel, Crystal A. Khan, Anne Rueger, et al. 2016. “Thermal Profiling Reveals Phenylalanine Hydroxylase as an off-Target of Panobinostat.” Nature Chemical Biology 12 (11): 908–10.

Blomen, Vincent A., Peter Májek, Lucas T. Jae, Johannes W. Bigenzahn, Joppe Nieuwenhuis, Jacqueline Staring, Roberto Sacco, et al. 2015. “Gene Essentiality and Synthetic Lethality in Haploid Human Cells.” Science 350 (6264): 1092–96.

Branca, Rui M. M., Lukas M. Orre, Henrik J. Johansson, Viktor Granholm, Mikael Huss, Åsa Pérez-Bercoff, Jenny Forshed, Lukas Käll, and Janne Lehtiö. 2014. “HiRIEF LC-MS Enables Deep Proteome Coverage and Unbiased Proteogenomics.” Nature Methods 11 (1): 59–62.

Calvo, Sarah E., Karl R. Clauser, and Vamsi K. Mootha. 2016. “MitoCarta2.0: An Updated Inventory of Mammalian Mitochondrial Proteins.” Nucleic Acids Research 44 (D1): D1251–57.

Chen, Ruibing, Yun Liu, Hao Zhuang, Baicai Yang, Kaiwen Hei, Mingming Xiao, Chunyu Hou, et al. 2017. “Quantitative Proteomics Reveals That Long Non-Coding RNA MALAT1 Interacts with DBC1 to Regulate p53 Acetylation.” Nucleic Acids Research 45 (17): 9947–59.

Csárdi, Gábor, Alexander Franks, David S. Choi, Edoardo M. Airoldi, and D. Allan Drummond. 2015. “Accounting for Experimental Noise Reveals That mRNA Levels, Amplified by Post-Transcriptional Processes, Largely Determine Steady-State Protein Levels in Yeast.” PLoS Genetics 11 (5): e1005206.

Culhane, Aedín C., Jean Thioulouse, Guy Perrière, and Desmond G. Higgins. 2005. “MADE4: An R Package for Multivariate Analysis of Gene Expression Data.” Bioinformatics 21 (11): 2789–90.

Dimitrakopoulos, Lampros, Ioannis Prassas, Eleftherios P. Diamandis, Alexey Nesvizhskii, Thomas Kislinger, Jacob Jaffe, and Andrei Drabovich. 2016. “Proteogenomics: Opportunities and Caveats.” Clinical Chemistry 62 (4): 551–57.

Duan, Guangyou, Xun Li, and Maja Köhn. 2015. “The Human DEPhOsphorylation Database DEPOD: A 2015 Update.” Nucleic Acids Research 43 (Database issue): D531–35.

Ezkurdia, Iakes, Jose Manuel Rodriguez, Enrique Carrillo-de Santa Pau, Jesús Vázquez, Alfonso Valencia, and Michael L. Tress. 2015. “Most Highly Expressed Protein-Coding Genes Have a Single Dominant Isoform.” Journal of Proteome Research 14 (4): 1880–87.

Fagerberg, Linn, Björn M. Hallström, Per Oksvold, Caroline Kampf, Dijana Djureinovic, Jacob Odeberg, Masato Habuka, et al. 2014. “Analysis of the Human Tissue-Specific Expression by Genome-Wide Integration of Transcriptomics and Antibody-Based Proteomics.” Molecular & Cellular Proteomics: MCP 13 (2): 397–406.

Franks, Alexander, Edoardo Airoldi, and Nikolai Slavov. 2017. “Post-Transcriptional Regulation across Human Tissues.” PLoS Computational Biology 13 (5): e1005535.

Futreal, P. Andrew, Lachlan Coin, Mhairi Marshall, Thomas Down, Timothy Hubbard, Richard Wooster, Nazneen Rahman, and Michael R. Stratton. 2004. “A Census of Human Cancer Genes.” Nature Reviews. Cancer 4 (3): 177–83.

Gaudet, Pascale, Pierre-André Michel, Monique Zahn-Zabal, Aurore Britan, Isabelle Cusin, Marcin Domagalski, Paula D. Duek, et al. 2017. “The neXtProt Knowledgebase on Human Proteins: 2017 Update.” Nucleic Acids Research 45 (D1): D177–82.

Geiger, Tamar, Ana Velic, Boris Macek, Emma Lundberg, Caroline Kampf, Nagarjuna Nagaraj, Mathias Uhlen, Juergen Cox, and Matthias Mann. 2013. “Initial Quantitative Proteomic Map of 28 Mouse Tissues Using the SILAC Mouse.” Molecular & Cellular Proteomics: MCP 12 (6): 1709–22.

Gevaert, Kris, Marc Goethals, Lennart Martens, Jozef Van Damme, An Staes, Grégoire R. Thomas, and Joël Vandekerckhove. 2003. “Exploring Proteomes and Analyzing Protein Processing by Mass Spectrometric Identification of Sorted N-Terminal Peptides.” Nature Biotechnology 21 (5): 566–69.

GTEx Consortium. 2013. “The Genotype-Tissue Expression (GTEx) Project.” Nature Genetics 45 (6): 580–85.

Hahne, Hannes, Fiona Pachl, Benjamin Ruprecht, Stefan K. Maier, Susan Klaeger, Dominic Helm, Guillaume Médard, Matthias Wilm, Simone Lemeer, and Bernhard Kuster. 2013. “DMSO Enhances Electrospray Response, Boosting Sensitivity of Proteomic Experiments.” Nature Methods 10 (10): 989–91.

Hao, Yun, and Nicholas P. Tatonetti. 2016. “Predicting G Protein-Coupled Receptor Downstream Signaling by Tissue Expression.” Bioinformatics 32 (22): 3435–43.

Hart, Traver, Megha Chandrashekhar, Michael Aregger, Zachary Steinhart, Kevin R. Brown, Graham MacLeod, Monika Mis, et al. 2015. “High-Resolution CRISPR Screens Reveal Fitness Genes and Genotype-Specific Cancer Liabilities.” Cell 163 (6): 1515–26.

Kearse, Michael G., and Jeremy E. Wilusz. 2017. “Non-AUG Translation: A New Start for Protein Synthesis in Eukaryotes.” Genes & Development 31 (17): 1717–31.

Kim, Min-Sik, Sneha M. Pinto, Derese Getnet, Raja Sekhar Nirujogi, Srikanth S. Manda, Raghothama Chaerkady, Anil K. Madugundu, et al. 2014. “A Draft Map of the Human Proteome.” Nature 509 (7502): 575–81.

Kleifeld, Oded, Alain Doucet, Ulrich auf dem Keller, Anna Prudova, Oliver Schilling, Rajesh K. Kainthan, Amanda E. Starr, Leonard J. Foster, Jayachandran N. Kizhakkedathu, and Christopher M. Overall. 2010. “Isotopic Labeling of Terminal Amines in Complex Samples Identifies Protein N-Termini and Protease Cleavage Products.” Nature Biotechnology 28 (3): 281–88.

Kolesnikov, Nikolay, Emma Hastings, Maria Keays, Olga Melnichuk, Y. Amy Tang, Eleanor Williams, Miroslaw Dylag, et al. 2015. “ArrayExpress Update--Simplifying Data Submissions.” Nucleic Acids Research 43 (Database issue): D1113–16.

Kustatscher, Georg, Piotr Grabowski, and Juri Rappsilber. 2017. “Pervasive Coexpression of Spatially Proximal Genes Is Buffered at the Protein Level.” Molecular Systems Biology 13 (8): 937.

Lambert, Samuel A., Arttu Jolma, Laura F. Campitelli, Pratyush K. Das, Yimeng Yin, Mihai Albu, Xiaoting Chen, Jussi Taipale, Timothy R. Hughes, and Matthew T. Weirauch. 2018. “The Human Transcription Factors.” Cell 172 (4): 650–65.

Lee, Chien-Yun, Dongxue Wang, Mathias Wilhelm, Daniel Paul Zolg, Tobias Schmidt, Karsten Schnatbaum, Ulf Reimer, et al. 2018. “Mining the Human Tissue Proteome for Protein Citrullination.” Molecular & Cellular Proteomics: MCP, April. https://doi.org/10.1074/mcp.RA118.000696.

Liu, Yansheng, Andreas Beyer, and Ruedi Aebersold. 2016. “On the Dependency of Cellular Protein Levels on mRNA Abundance.” Cell 165 (3): 535–50.

Marx, Harald, Hannes Hahne, Susanne E. Ulbrich, Angelika Schnieke, Oswald Rottmann, Dmitrij Frishman, and Bernhard Kuster. 2017. “Annotation of the Domestic Pig Genome by Quantitative Proteogenomics.” Journal of Proteome Research 16 (8): 2887–98.

Matthews, Dwight E. 2007. “An Overview of Phenylalanine and Tyrosine Kinetics in Humans.” The Journal of Nutrition 137 (6 Suppl 1): 1549S–1555S; discussion 1573S–1575S.

Mertins, Philipp, D. R. Mani, Kelly V. Ruggles, Michael A. Gillette, Karl R. Clauser, Pei Wang, Xianlong Wang, et al. 2016. “Proteogenomics Connects Somatic Mutations to Signalling in Breast Cancer.” Nature 534 (7605): 55–62.

Na, Chan Hyun, Mustafa A. Barbhuiya, Min-Sik Kim, Steven Verbruggen, Stephen M. Eacker, Olga Pletnikova, Juan C. Troncoso, et al. 2018. “Discovery of Noncanonical Translation Initiation Sites through Mass Spectrometric Analysis of Protein N Termini.” Genome Research 28 (1): 25–36.

Nesvizhskii, Alexey I. 2014. “Proteogenomics: Concepts, Applications and Computational Strategies.” Nature Methods 11 (11): 1114–25.

Omenn, Gilbert S., Lydie Lane, Emma K. Lundberg, Christopher M. Overall, and Eric W. Deutsch. 2017. “Progress on the HUPO Draft Human Proteome: 2017 Metrics of the Human Proteome Project.” Journal of Proteome Research, October. https://doi.org/10.1021/acs.jproteome.7b00375.

Rappsilber, Juri, Matthias Mann, and Yasushi Ishihama. 2007. “Protocol for Micro-Purification, Enrichment, Pre-Fractionation and Storage of Peptides for Proteomics Using StageTips.” Nature Protocols 2 (8): 1896–1906.

Ruprecht, Benjamin, Dongxue Wang, Riccardo Zenezini Chiozzi, Li-Hua Li, Hannes Hahne, and Bernhard Kuster. 2017. “Hydrophilic Strong Anion Exchange (hSAX) Chromatography Enables Deep Fractionation of Tissue Proteomes.” Methods in Molecular Biology 1550: 69–82.

Schmidt, Tobias, Patroklos Samaras, Martin Frejno, Siegfried Gessulat, Maximilian Barnert, Harald Kienegger, Helmut Krcmar, et al. 2018. “ProteomicsDB.” Nucleic Acids Research 46 (D1): D1271–81.

Schwanhäusser, Björn, Dorothea Busse, Na Li, Gunnar Dittmar, Johannes Schuchhardt, Jana Wolf, Wei Chen, and Matthias Selbach. 2011. “Global Quantification of Mammalian Gene Expression Control.” Nature 473 (7347): 337–42.

Thul, Peter J., Lovisa Åkesson, Mikaela Wiking, Diana Mahdessian, Aikaterini Geladaki, Hammou Ait Blal, Tove Alm, et al. 2017. “A Subcellular Map of the Human Proteome.” Science 356 (6340). https://doi.org/10.1126/science.aal3321.

Toprak, Umut H., Ludovic C. Gillet, Alessio Maiolica, Pedro Navarro, Alexander Leitner, and Ruedi Aebersold. 2014. “Conserved Peptide Fragmentation as a Benchmarking Tool for Mass Spectrometers and a Discriminating Feature for Targeted Proteomics.” Molecular & Cellular Proteomics: MCP 13 (8): 2056–71.

Trapnell, Cole, Brian A. Williams, Geo Pertea, Ali Mortazavi, Gordon Kwan, Marijke J. van Baren, Steven L. Salzberg, Barbara J. Wold, and Lior Pachter. 2010. “Transcript Assembly and Quantification by RNA-Seq Reveals Unannotated Transcripts and Isoform Switching during Cell Differentiation.” Nature Biotechnology 28 (5): 511–15.

Uhlén, Mathias, Linn Fagerberg, Björn M. Hallström, Cecilia Lindskog, Per Oksvold, Adil Mardinoglu, Åsa Sivertsson, et al. 2015. “Proteomics. Tissue-Based Map of the Human Proteome.” Science 347 (6220): 1260419.

Uhlén, Mathias, Björn M. Hallström, Cecilia Lindskog, Adil Mardinoglu, Fredrik Pontén, and Jens Nielsen. 2016. “Transcriptomics Resources of Human Tissues and Organs.” Molecular Systems Biology 12 (4): 862.

Vizcaíno, Juan A., Eric W. Deutsch, Rui Wang, Attila Csordas, Florian Reisinger, Daniel Ríos, José A. Dianes, et al. 2014. “ProteomeXchange Provides Globally Coordinated Proteomics Data Submission and Dissemination.” Nature Biotechnology 32 (3): 223–26.

Vogel, Christine, Raquel de Sousa Abreu, Daijin Ko, Shu-Yun Le, Bruce A. Shapiro, Suzanne C. Burns, Devraj Sandhu, Daniel R. Boutz, Edward M. Marcotte, and Luiz O. Penalva. 2010. “Sequence Signatures and mRNA Concentration Can Explain Two-Thirds of Protein Abundance Variation in a Human Cell Line.” Molecular Systems Biology 6 (August): 400.

Wang, Tim, Kivanç Birsoy, Nicholas W. Hughes, Kevin M. Krupczak, Yorick Post, Jenny J. Wei, Eric S. Lander, and David M. Sabatini. 2015. “Identification and Characterization of Essential Genes in the Human Genome.” Science 350 (6264): 1096–1101.

Wang, Xiaojing, and Bing Zhang. 2013. “customProDB: An R Package to Generate Customized Protein Databases from RNA-Seq Data for Proteomics Search.” Bioinformatics 29 (24): 3235–37.

Wilhelm, Mathias, Hannes Hahne, Mikhail Savitski, Harald Marx, Simone Lemeer, Marcus Bantscheff, and Bernhard Kuster. 2017. “Wilhelm et Al. Reply.” Nature 547 (7664): E23.

Wilhelm, Mathias, Judith Schlegl, Hannes Hahne, Amin Moghaddas Gholami, Marcus Lieberenz, Mikhail M. Savitski, Emanuel Ziegler, et al. 2014. “Mass-Spectrometry-Based Draft of the Human Proteome.” Nature 509 (7502): 582–87.

Wishart, David S., Yannick D. Feunang, An C. Guo, Elvis J. Lo, Ana Marcu, Jason R. Grant, Tanvir Sajed, et al. 2018. “DrugBank 5.0: A Major Update to the DrugBank Database for 2018.” Nucleic Acids Research 46 (D1): D1074–82.

Yu, Guangchuang, Fei Li, Yide Qin, Xiaochen Bo, Yibo Wu, and Shengqi Wang. 2010. “GOSemSim: An R Package for Measuring Semantic Similarity among GO Terms and Gene Products.” Bioinformatics 26 (7): 976–78.

Yu, Guangchuang, Li-Gen Wang, Yanyan Han, and Qing-Yu He. 2012. “clusterProfiler: An R Package for Comparing Biological Themes among Gene Clusters.” Omics: A Journal of Integrative Biology 16 (5): 284–87.

Zhang, Bing, Jing Wang, Xiaojing Wang, Jing Zhu, Qi Liu, Zhiao Shi, Matthew C. Chambers, et al. 2014. “Proteogenomic Characterization of Human Colon and Rectal Cancer.” Nature 513 (7518): 382–87.

Zolg, Daniel P., Mathias Wilhelm, Karsten Schnatbaum, Johannes Zerweck, Tobias Knaute, Bernard Delanghe, Derek J. Bailey, et al. 2017. “Building ProteomeTools Based on a Complete Synthetic Human Proteome.” Nature Methods 14 (3): 259–62.

